# RSL24D1 sustains steady-state ribosome biogenesis and pluripotency translational programs in embryonic stem cells

**DOI:** 10.1101/2021.05.27.443845

**Authors:** Sébastien Durand, Marion Bruelle, Fleur Bourdelais, Bigitha Bennychen, Juliana Blin-Gonthier, Caroline Isaac, Aurélia Huyghe, Antoine Seyve, Christophe Vanbelle, David Meyronet, Frédéric Catez, Jean-Jacques Diaz, Fabrice Lavial, Emiliano P. Ricci, François Ducray, Mathieu Gabut

## Abstract

Embryonic stem cell (ESC) fate decisions are regulated by a complex molecular circuitry that requires tight and coordinated gene expression regulations at multiple levels from chromatin organization to mRNA processing. Recently, ribosome biogenesis and translation have emerged as key regulatory pathways that efficiently control stem cell homeostasis. However, the molecular mechanisms underlying the regulation of these pathways remain largely unknown to date. Here, we analyzed the expression, in mouse ESCs, of over 300 genes involved in ribosome biogenesis and we identified RSL24D1 as the most differentially expressed between self-renewing and differentiated ESCs. RSL24D1 is highly expressed in multiple mouse pluripotent stem cell models and its expression profile is conserved in human ESCs. RSL24D1 is associated with nuclear pre-ribosomes and is required for the maturation and the synthesis of 60S subunits in mouse ESCs. Interestingly, RSL24D1 depletion significantly impairs global translation, particularly of key pluripotency factors, including POU5F1 and NANOG, as well as components of the polycomb repressive complex 2 (PRC2). Consistently, RSL24D1 is required for mouse ESC self-renewal and proliferation. Taken together, we show that RSL24D1-dependant ribosome biogenesis is required to both sustain the expression of pluripotent transcriptional programs and silence developmental programs, which concertedly dictate ESC homeostasis.

## Introduction

Pluripotent stem cells (PSCs), including embryonic stem cells (ESCs) and induced pluripotent stem cells (iPSCs), have the unique abilities to self-renew in a naive state while remaining competent to differentiate into a spectrum of lineages that compose developing embryonic tissues. This ambivalent state is tightly and dynamically coordinated at different steps of gene expression, which have been extensively described at the chromatin (1), transcriptional (2, 3) and post-transcriptional levels (4-6). This led to the identification of key regulatory epigenetic and transcriptional programs that rapidly rewire gene expression and regulate complex cell fate transitions in response to environmental cues (1, 3, 7, 8). For instance, chromatin modifications mediated by the Polycomb Repressive Complexes 1 and 2 (PRC1 and PRC2) have been shown to play pivotal roles in the transcriptional regulation of pluripotency and differentiation of PSCs (1, 7-9). However, multiple studies suggest that, in many circumstances, messenger RNA (mRNA) and protein levels are poorly correlated, including in ESCs and iPSCs, therefore highlighting the importance of translational regulations for shaping the cellular proteome landscape required for cell fate changes (10-15). The relevance of such observations is strengthened by recent studies highlighting that translation is regulated during PSCs differentiation, with a lower translation efficiency in undifferentiated ESCs, but also in adult stem cell models compared to differentiated progenies (16-19). Remarkably, murine ESC proliferation is regulated by interdependent layers involving translation, euchromatin organization and transcriptional control (20). This is well illustrated by HTATSF1-dependent coordination of protein synthesis with ribosomal RNA (rRNA) processing during human ESC differentiation (21). Taken together, this compelling evidence therefore demonstrates that the regulation of protein synthesis plays a key role in defining stem cell fate.

In eukaryotes, ribosome biogenesis is a complex and multistep process that involves different cellular compartments, consecutively the nucleoli, the nucleoplasm and the cytoplasm, as well as over 280 ribosome biogenesis factors (RBFs) and different families of non-coding RNAs (22-24). Many studies have established that ribosome biogenesis is finely regulated in stem cells and may directly control stem cell properties (16). First, despite having a low protein synthesis activity, ESCs display higher levels of rRNA transcription rates compared to endodermal-lineage committed cells (25). Moreover, the nuclear remodelling complex NoRC has been shown to coordinate rRNA transcription and proliferation rates in mouse ESCs (26). Similarly, several RFBs are expressed at higher levels in ESCs compared to differentiated progenies (27), including Nucleolin (28), Dyskerin Pseudouridine Synthase 1 (DKC1) (29) and Fibrillarin (FBL) (30), and are required for ESC self-renewal. In addition to these well-described essential RBFs controlling rRNA expression and post-transcriptional modifications, it appears that ribosome subunit-specific RBFs are required to support ESC maintenance. Indeed, several factors implicated in the maturation of the 40S small ribosome subunit (SSU) are preferentially expressed in naive ESCs compared to differentiated progenies and support the translation of key pluripotency transcription factors (PTFs) such as NANOG (31). Notably, Notchless, a RBF of the 60S large ribosome subunit (LSU), is required for the Inner Cell Mass survival during early mouse embryogenesis, yet it is unclear whether ribosome biogenesis is implicated in this context, in contrast to Notchless functions in adult stem cells homeostasis (32-35). Thus, to date, the contributions of ribosome subunit-specific RBFs, especially of pre-60S RBFs, to the steady-state stoichiometry of the 40S and 60S subunits and their impact on the regulation of PSC fate decision remain unclear and should be further investigated.

Here, we show that RSL24D1, a conserved homolog of the yeast pre-60S maturation and export factor Rlp24, is expressed at high levels in mouse and human PSCs compared to differentiated progenies. We demonstrate that RSL24D1 is essential for the maturation of the LSU in pluripotent mouse ESCs. RSL24D1 is also required to maintain a steady-state level of translation, in particular of PTFs, such as NANOG and POU5F1 that control pluripotent transcriptional programs, but also of PRC2 factors that maintain repressive H3K27me3 marks over developmental and differentiation genes to prevent their premature activation. Moreover, high levels of RSL24D1 are required to support mouse ESC proliferation and self-renewal. Altogether, these results establish for the first time that a *bona fide* 60S biogenesis and the resulting proper translation status are coordinated with transcription and chromatin regulation networks to control ESC homeostasis.

## Results

### Rsl24d1 expression is enriched in murine and human pluripotent cells

To identify factors contributing to ribosome assembly in pluripotent stem cells, we first defined the mRNA expression profiles of 303 genes, including RBFs and ribosomal proteins (RPs), in murine iPSC clones, ESCs and differentiated cell lines using publicly available RNA-seq data (36) (Supplemental Fig. S1A and Table 1). The majority (70%) of factors associated with the biogenesis of the LSU and SSU, as well as RPs were expressed at higher levels in pluripotent cells compared to differentiated cells (fold change >1,5) (Fig. 1A). Among these factors, we identified Rsl24d1, a predicted ribosome biogenesis protein, which displayed the most striking expression change between differentiated mouse cell lines and PSCs (Log2 fold change > 13,3) (Fig. 1A). We next confirmed that RSL24D1 was also expressed at a higher level in mouse pluripotent CGR8 ESCs cultured either in serum+LIF (ESC^FBS^) or in 2i-induced naïve ground state (ESC^2i^) conditions compared to *in vitro* ESC-derived 12-day old differentiated embryoid bodies (EBs) containing beating cardiomyocytes (EB^12^) (Fig. 1B). In contrast, the expression of RPL8, a canonical RP of the LSU, remained globally unchanged at the protein level as ESCs differentiate into EBs (Fig. 1B).

**Figure 1.**
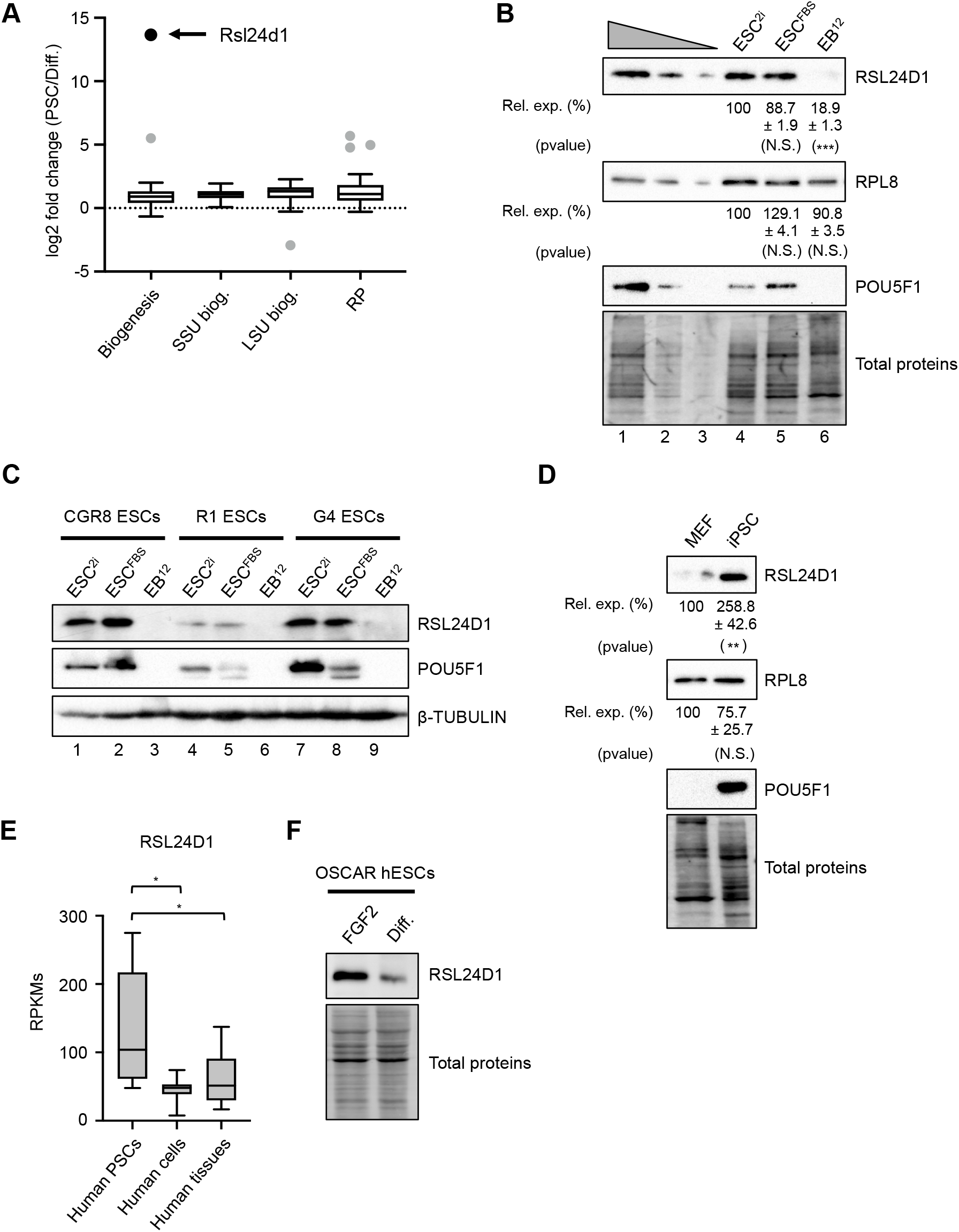
Rsl24d1 expression is enriched in PSCs. (A) Box plot representation of mRNA expression changes in mouse PSCs (5 ESC lines and 2 iPSC clones) and 6 differentiated cell lines, measured by RNA-seq, for 4 functional gene categories: factors involved in common biogenesis steps (Biogenesis), specific biogenesis factors of the 40S (SSU biog.) or of the 60S subunits (LSU biog.), and ribosomal proteins (RP). Outlier values are represented by individual grey circles and Rsl24d1 is indicated by a black circle. Additional details are provided in Figure S1A and individual data are available in Table S1. (B) Representative western blot analyses of RSL24D1 (n=6), POU5F1 (n=4) and RPL8 (n=5) in mouse CGR8 ESCs cultivated in ground state (ESC^2i^) or naive (ESC^FBS^) conditions or differentiated in embryoid bodies (EB) for 12 days (EB^12^). Lanes 1 to 3 correspond to serial of dilutions of ESC^FBS^ (1:1, 1:3 and 1:9, respectively). Tri-Chloro-Ethanol (TCE) labeling of tryptophan-containing proteins (referred to as Total Proteins) is used for normalization. Quantifications of RSL24D1 and RPL8 signals normalized to total proteins and relative to the ESC^2i^ condition (Rel. exp. (%)) are indicated below each corresponding panels. Indicated P values are relative to ESC^FBS^ conditions (paired two-tailed Student’s t-test): *** pval<0.001, N.S.: not significant (pval>0.05). (C) Representative western blots of RSL24D1 and POU5F1 in 3 unrelated mouse ESC lines (CGR8, R1 and G4) cultivated in ESC^2i^, ESC^FBS^ and differentiated (EB^12^) conditions, as in panel B. β-TUBULIN levels are shown as a loading control. (D) Representative western blot analyses of RSL24D1 (n=6), POU5F1 (n=4) and RPL8 (n=4) in MEFs and in iPSCs reprogrammed from MEFs derived from doxycycline-inducible Col1a1-tetO-OKMS mice after ectopic induction of Pou5f1, Sox2, Klf4 and cMyc expression. TCE labeling of tryptophan-containing proteins (referred as Total Proteins) is used for normalization. Quantifications of RSL24D1 and RPL8 signals normalized to total protein and relative to the “MEF” conditions (Rel. exp. (%)) are indicated below each corresponding panels. Indicated P values are relative to “MEF” conditions (paired two-tailed Student’s t-test): ** pval<0.005, N.S.: not significant (pval>0.05). (E) Box plot representation of normalized RSL24D1 mRNA expression levels in RPKMs (Reads per kilo base per million mapped reads) across 5 human PSC models, 7 human cell lines and 16 adult tissues based on published RNA-seq profiles (36). P values are indicated and are relative to human PSCs (unpaired two-tailed Student’s t-test): * pval<0,05. (F) Representative western blots of RSL24D1 in human OSCAR ESCs maintained in pluripotent state in the presence of FGF2 or *in vitro* differentiated into EBs (Diff.). TCE labeling of proteins (“Total Proteins”) is used as a loading control.

To get more insights into the regulation of Rsl24d1 expression in PSCs, we next assessed the dynamics of Rsl24d1 expression during the kinetics of ESC-derived EB differentiation. RT-qPCR assays confirmed that the expression of pluripotency transcription factors Pou5f1, Klf4, Nanog and Sox2 rapidly decreased upon ESC differentiation into EBs (Supplemental Fig. S1B). Interestingly, Rsl24d1 is expressed at the highest level in mouse CGR8 ESCs and is progressively downregulated after differentiation initiation to reach its lowest expression level in EB^12^, thereby correlating with the expression profiles of key PTFs. Consistent with mRNA levels, RSL24D1 protein expression was strongly lowered after 5 days of EB formation and further decreased as the differentiation proceeded to reach a minimal expression in EB^12^ (Supplemental Fig. S1C). The expression of additional RPs also decreased, yet to a lower extent than RSL24D1, while the downregulation of two biogenesis factors, EIF6 and NOG1, rather followed RSL24D1’s profile. In addition, we assessed the expression of RSL24D1 in two additional mouse ESCs lines (R1 and G4) cultured in similar conditions (ESC^2i^, ESC^FBS^, EB^12^) (Fig. 1C). Although RSL24D1 was expressed at different basal levels in the three ESC models, these results confirmed that RSL24D1 levels were significantly higher in ESCs maintained in pluripotent states than in ESC-derived differentiated EBs. Altogether, these results convincingly demonstrate that RSL24D1 expression is high in murine ESCs and strongly decreases upon differentiation.

We then hypothesized that the expression of Rsl24d1 would rather be determined by the pluripotency status than by the embryonic origin. Therefore, pluripotent cells from non-embryonic origin should also express high levels of RSL24D1 compared to their differentiated counterparts. To evaluate this hypothesis, iPSCs were generated after somatic reprogramming of mouse embryonic fibroblasts (MEFs) by forced expression of Pou5f1, Klf4, c-Myc and Sox2 (OKMS) (37). The kinetics of somatic reprogramming was assessed by the activation of both endogenous Pou5f1 and Nanog mRNAs (Supplemental Fig. S1D). While RPL8 expression remained globally unchanged, RSL24D1 expression was highly increased in 14-day old iPSCs compared to parental MEFs (Fig. 1D). Since early steps of iPSC formation are highly heterogeneous and stochastic (38, 39), we next investigated Rsl24d1 expression in cells with enhanced reprogramming potential at the single cell level from published data (40). Interestingly, single-cell RNA-seq data and fate trajectory detection by Guo and colleagues revealed that Rsl24d1 expression was strongly enhanced in a continuum of cells representing different stages of active reprogramming (pre-PCs) compared to cells that are engaged in earlier steps of the reprogramming path (RP) (Supplemental Fig. S1E). Strikingly, Rsl24d1 levels were even further increased in chimera-competent reprogrammed cells (PCs) expressing high levels of pluripotency factors including Nanog and Esrrb (Supplemental Fig. S1E). Altogether, these observations suggest that Rsl24d1 expression is significantly enriched in mouse PSCs regardless of their embryonic origins.

We next investigated whether the regulation of Rsl24d1 expression was evolutionarily conserved in human PSCs. RNA-seq analysis indicated that, similarly to their murine counterparts, human PSCs expressed higher levels of *RSL24D1* mRNAs compared to differentiated cell lines or tissues (Fig. 1E) (36). Interestingly, western blot analyses of RSL24D1 in human OSCAR ESCs cultured in FGF2-supplemented self-renewal media and in ESC-derived EBs (“Diff.”) revealed a marked decrease of RSL24D1 upon differentiation (Fig. 1F) (41). In addition, RSL24D1 was expressed at similar levels in both human OSCAR and H9 ESCs maintained either in primed (TL and FGF2) or naive-like (TL2i) conditions (41) (Supplemental Fig. S1F), whereas RSL24D1 expression was rather low in adult human tissues compared to human ESCs (Supplemental Fig. S1G). Taken together, these results indicate that Rsl24d1 expression is high in PSCs and decreases upon loss of pluripotency, and that this regulation is evolutionarily conserved.

### RSL24D1 is a biogenesis factor associated with nuclear pre-ribosomes in mouse ESCs

The role of RSL24D1 in higher eukaryotes remains unknown, therefore, to gain insight into its molecular functions in mouse ESCs, we first compared its sequence and structure with conserved homologs. Indeed, Rlp24, the yeast homolog of RSL24D1, is a ribosome biogenesis factor involved in the export of nuclear pre-60S ribosomal particles from the nucleus to the cytoplasm where they undergo the final steps of maturation, including Rlp24 substitution by the canonical ribosomal protein Rpl24 (23, 42). Multiple protein alignments of yeast Rlp24 to higher eukaryote homologs, including murine and human RSL24D1, revealed that the first 130 amino acids of Rlp24 were well conserved from yeast to human (Supplemental Fig. S2A, B). In particular, RSL24D1 homologs were strongly conserved in mammals, with a sequence identity over 96%, but lacked the yeast-specific C-terminal extension of Rlp24.

To further assess the degree of conservation between higher eukaryote RSL24D1 and its yeast homolog, we next compared the structure of mouse RSL24D1 with cryo-EM structures of yeast and human protein homologs, which have been recently obtained from nuclear pre-60S intermediates (Fig. 2A) (43-46). Similarly to yeast, cryo-EM structures of human pre-60S particles revealed the presence of RSL24D1 in nucleoplasmic stages of pre-60S assembly (states pre-A and A) while it was absent from later cytoplasmic stages of pre-60S maturation (46). As the N-terminal region of Rlp24 homologs was the most conserved during evolution, we used these cryo-EM structures to model the structure of the mouse RSL24D1 N-terminus from its amino acid sequence. Interestingly, structure alignments indicate that the predicted structure of the first 135 amino acids of mouse RSL24D1 almost perfectly matches the yeast Rlp24 and human RSL24D1 structures from nuclear pre-60S intermediates (Fig. 2A) (46). Altogether, these results strongly support a conserved function of RSL24D1 in the nuclear maturation of the pre-60S particles in mouse ESCs.

**Figure 2.**
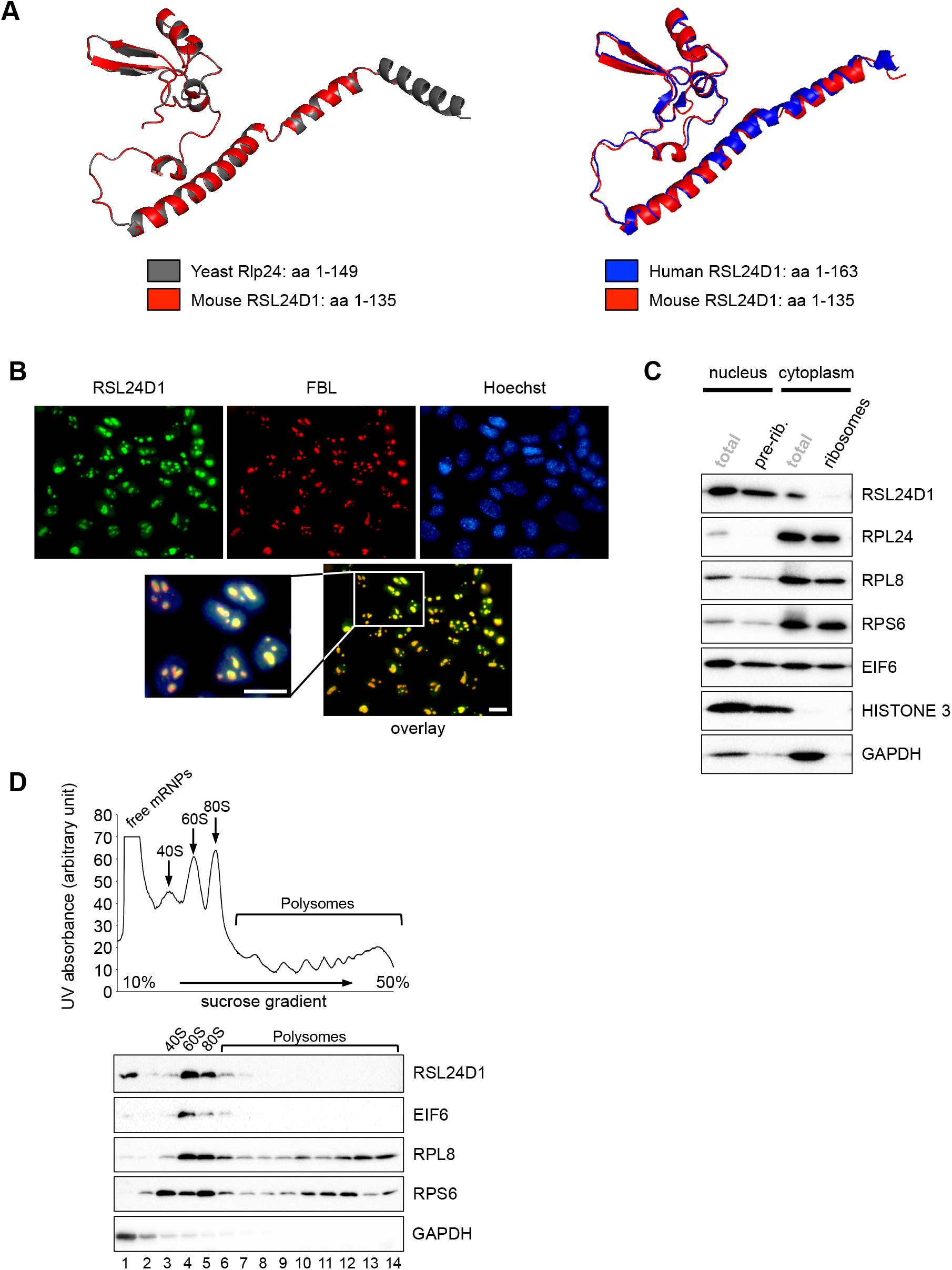
RSL24D1 protein is associated with pre-60S subunits in mouse ESCs. (A) Comparison of the mouse RSL24D1 predicted structure (amino acids 1-135, red colour) with the yeast Rlp24 structure (amino acids 1-149, PDB 6N8J grey colour, left panel) and the human RSL24D1 structure (amino acids 1-163, PDB 6LSS, blue colour, right panel) (45-46). The mouse RSL24D1 protein structure was predicted from the 6N8J and 6LSS structures using the swiss-model structure assessment tool (87). (B) Representative images of naive CGR8 cells stained with Hoechst and with anti-FBL and anti-RSL24D1 antibodies (20X objective). (C) Nucleo-cytoplasmic fractionations (“total”) followed by sucrose cushion purifications of nuclear pre-ribosomes (“pre-rib.”) and cytoplasmic ribosomes (“ribosomes”) from CGR8 ESC^FBS^. Representative western blot analysis of RSL24D1, RPL24 and RPL8 (LSU), RPS6 (SSU) and EIF6 (RBF). HISTONE H3 and GAPDH are shown as specific nuclear and cytoplasmic proteins, respectively. (D) Polysome profiling by centrifugation on sucrose gradient of CGR8 ESC^FBS^ cytoplasmic extracts. Ribosome-free fractions (free mRNPs), 40S, 60S, 80S monosomes and polysomes are detected by UV-absorbance and indicated on the absorbance curve (upper panel). Representative western blots of gradient fractions using antibodies targeting RSL24D1, EIF6 (LSU), RPL8 (LSU) and RPS6 (SSU) (lower panel). GAPDH is used as a control of the free mRNPs. The fractions are indicated below the GAPDH panel.

To test this hypothesis, we next analyzed the localization of RSL24D1 in mouse CGR8 ESCs. Since no difference in RSL24D1 expression was observed between the ESC^2i^ and ESC^FBS^ conditions, all following experiments were performed with CGR8 cells cultured in media containing serum and LIF (ESC^FBS^). In these conditions, RSL24D1 was expressed in all colony-forming ESCs regardless of POU5F1 steady-state levels (Supplemental Fig. S2C) and was predominantly concentrated within nuclear foci containing FBL (Fig. 2B), therefore suggesting that RSL24D1 is mostly located in ESC nucleoli (47).

We then asked whether RSL24D1 was associated with pre-ribosomal particles in mouse ESC^FBS^ colonies. Following cell fractionation, pre-ribosomal and ribosomal particles were respectively isolated from nuclear and cytoplasmic fractions by ultracentrifugation on sucrose cushions. In contrast to RPL8 and RPS6 that are both present in nuclear pre-ribosomes and in cytoplasmic ribosomes, RPL24 is exclusively present in cytoplasmic ribosomes (Fig. 2C). This is consistent with observations that Rpl24 replaces Rlp24 in pre-60S particles after nuclear export in yeast and human (23, 42, 46). In addition, the LSU biogenesis factor EIF6, homolog of yeast Tif6, co-purifies with both nuclear and cytoplasmic ribosomal particles, whereas RSL24D1 is predominantly detected in nuclear pre-ribosomes (Fig. 2C). Cell fractionation assays confirmed that RSL24D1 is predominantly detected in the nuclear fraction (68%) and to a lower extent in the cytoplasmic fraction (32%), while RPL24 is almost exclusively present in the cytoplasm (95%) (Supplemental Fig. S2D). RSL24D1 is also slightly associated with cytoplasmic ribosomes yet to a lower extent compared to EIF6, suggesting that RSL24D1 is rapidly removed from cytoplasmic pre-ribosomes and most likely replaced by RPL24 after nuclear export. To confirm that RSL24D1 is associated with pre-60S particles, cytoplasmic fractions were analyzed by polysome profiling assays to separate the 40S (SSU), 60S (LSU), 80S (monosomes) and polysomes (Supplemental Fig. S2E). As expected, RPL8 and RPS6 are predominantly detected in 60S and 40S ribosomal fractions, respectively, and both proteins are also present in 80S monosomes and polysomes (Fig. 2D). In contrast, RSL24D1 and EIF6 are mainly detected in 60S fractions (lane 4). RSL24D1 and EIF6 are also found at a lower extent in fractions enriched in monosomes (lanes 5 and 6), most likely due to an incomplete separation of 80S fractions from 60S fractions, while they are not significantly detected in the 40S fractions (lane 3). Altogether, these results suggest for the first time that RSL24D1 is a ribosome biogenesis factor in higher eukaryotes, which is incorporated into nucleolar pre-60S particles, transits to the cytoplasm and is subsequently removed from cytoplasmic pre-60S.

### RSL24D1 depletion impairs both ribosome biogenesis and translation

To confirm that RSL24D1 is involved in ribosome biogenesis, we then determined whether its depletion impacts the accumulation and activity of mature cytoplasmic ribosomes. Rsl24d1 siRNA treatment for 72 hours resulted in an efficient depletion of RSL24D1 (> 67%) in mouse ESCs compared to control non-targeting siRNAs (Fig. 3A and Supplemental Fig. 3A). Interestingly, RSL24D1 depletion did not detectably affect the structure and the number of FBL-containing nucleoli (data not shown), therefore suggesting that decreasing RSL24D1 levels did not induce major disruptions of early ribosome biogenesis (48).

**Figure 3.**
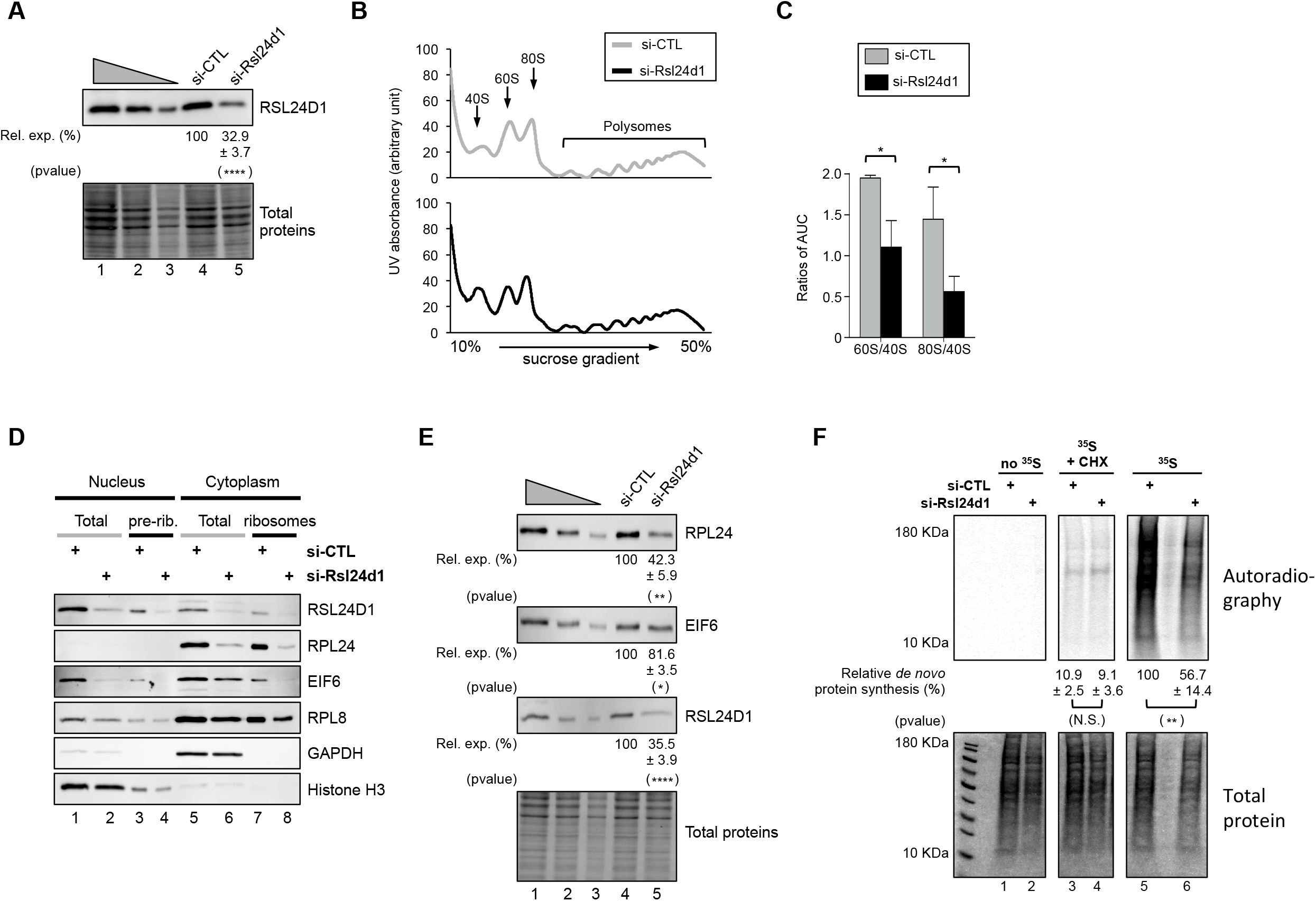
RSL24D1 depletion alters ribosome biogenesis and protein translation. (A) Representative immunoblot of RSL24D1 in total extracts from si-CTL and si-Rsl24d1 treated CGR8 ESC^FBS^. Lanes 1 to 3 correspond to serial dilutions of ESC^FBS^ (1:1, 1:3 and 1:9, respectively). TCE labeling of tryptophan-containing proteins (referred as Total Proteins) is used for normalization. Quantifications of RSL24D1 signals normalized to total proteins and relative to the si-CTL-treated conditions (Rel. exp. (%)) are indicated below each panel. The indicated P value is relative to the si-CTL-treated conditions (unpaired two-tailed Student’s t test): **** pval<0.0001. (B) Polysome profiling by centrifugation on sucrose gradient of cytoplasmic extracts from naive CGR8 cells treated with non-targeting siRNAs (grey color) or siRNAs targeting Rsl24d1 (black color). 40S, 60S, 80S monosome and polysomes are detected by UV-absorbance and indicated on the absorbance curve. (C) Histograms indicating ratios of 60S/40S and 80S/40S absorbance peaks calculated by determining the area under the curve (AUC) for the 40S, 60S and 80S absorbance signals (n=3). P values are indicated (unpaired two-tailed Student’s t test): * (pval<0.05). (D) Nucleo-cytoplasmic fractionation followed by ribosome purifications on sucrose cushions (“pre-rib.” and “ribosomes”) of CGR8 ESC^FBS^ transfected with non-targeting (si-CTL) or Rsl24d1-targeting siRNAs. Nuclear and cytoplasmic extracts are both indicated by “Total”. Western blots are probed with anti-RSL24D1, -RPL24, -RPL8 and -EIF6 antibodies. Immunoblots for HISTONE H3 and GAPDH are shown as specific nuclear and cytoplasmic proteins, respectively. (E) Representative immunoblots of RSL24D1, RPL24 and EIF6 (n=3) in total extracts from si-CTL and si-Rsl24d1 treated CGR8 ESC^FBS^. Lanes 1 to 3 correspond to serial dilutions of si-CTL-treated ESC^FBS^ (1:1, 1:3 and 1:9, respectively). TCE labeling of tryptophan-containing proteins (referred as Total Proteins) is used for normalization. Quantifications of the RPL24, EIF6 and RSL24D1 signals normalized to total proteins and relative to the si-CTL-treated condition (Rel. exp. (%)) are indicated below each panel. The indicated P values are relative to the si-CTL-treated conditions (unpaired two-tailed Student’s t test): **** pval<0.0001, ** pval<0.01, * pval<0.05. (F) Representative autoradiography (upper panel) and coomassie staining (lower panel) of SDS-PAGE of total extracts from si-CTL and si-Rsl24d1 treated CGR8 ESC^FBS^, in the presence or absence of Cycloheximide (CHX) and ^35^S-labelled methionine and cysteine. Quantifications of ^35^S signals (autoradiography) are normalized to total proteins (coomassie staining) and expressed relative to si-CTL- and ^35^S-methionine-treated ESC^FBS^ (n=3). Indicated P values are calculated relative to si-CTL-treated conditions (unpaired two-tailed Student’s t test): ** pval<0.01, N.S.: pval>0.05.

Next, the effect of RSL24D1 depletion on ribosome production was assessed by polysome profiling assays conducted on control- or Rsl24d1-siRNA treated ESCs. The transient depletion of RSL24D1 significantly imbalanced the accumulation of 40S, 60S and 80S particles (Fig. 3B), notably causing a major loss of 60S and 80S relative to 40S subunits (Fig. 3C). To achieve a more efficient and stable depletion of RSL24D1, we designed two independent shRNAs targeting Rsl24d1 mRNAs (sh-Rsl24d1-1 and sh-Rsl24d1-2), which resulted in a 51% and 93% depletion of RSL24D1, respectively, compared to control shRNAs (Supplemental Fig. S3B). Interestingly, expression of sh-Rsl24d1-2, which provided the most robust silencing of RSL24D1, also caused an impaired accumulation of the 80S and 60S subunits, to a similar extent than siRNA-treated cells (Supplemental Fig. S3C and S3D).

In order to further characterize molecular alterations resulting from RSL24D1 depletions, nuclear and cytoplasmic ribosomal fractions were isolated from ESCs treated with control- and Rsl24d1-targetting siRNAs (Fig. 3D). The depletion of RSL24D1 resulted in a significant loss of RSL24D1 and EIF6 association to nuclear pre-ribosomes (lanes 3-4) and to cytoplasmic particles (lanes 7-8). RPL24 inclusion into the cytoplasmic ribosomal particles was also strongly impaired upon RSL24D1 knockdown (lanes 7-8), while the presence of the canonical RPL8 in both nuclear pre-ribosomes and cytoplasmic ribosomes was not significantly affected (lanes 1-8). Moreover, we observed a consistent decrease of RPL24 and EIF6 expression upon RSL24D1 depletion in total cytoplasmic (lanes 5-6) and nuclear (lanes 1-2) fractions, respectively. Western blot assays on total extracts confirmed that RPL24 and EIF6 expression levels decreased upon RSL24D1 knockdown (Fig. 3E). These observations suggest that defects in EIF6 and RPL24 association with pre-60S particles might interfere with the stability of these proteins. Altogether, these results indicate that RSL24D1 is required for the *bona fide* maturation of pre-60S particles, at least by allowing EIF6 association to nuclear pre-ribosomes and RPL24 recruitment to mature cytoplasmic ribosomes. Hence, RSL24D1 plays a critical role in the production of mature 60S in mouse ESCs.

Finally, as RSL24D1 depletion impacts the LSU production, we hypothesized that RSL24D1 knockdown may affect global protein synthesis in ESCs. ^35^S pulse-chase labelling assays were performed on CGR8 cells transfected with si-CTL or si-Rsl24d1, in the presence or absence of cycloheximide (CHX), an inhibitor of translation elongation (Fig. 3F). As expected, CHX significantly prevented the incorporation of both ^35^S -labeled methionine and cysteine in newly synthetized proteins (lanes 3-4). Strikingly, RSL24D1 depletion caused a significant reduction (43%) of *de novo* protein synthesis in ESCs (lanes 5-6), therefore demonstrating that RSL24D1 loss decreases the global protein synthesis activity of ESCs. Altogether, these results strongly suggest that RSL24D1 is as an essential ribosome biogenesis factor of the LSU, which is required for the steady-state protein synthesis in pluripotent ESCs.

### Rsl24d1 is required to maintain pluripotent transcriptional programs

We next hypothesized that the effects of Rsl24d1 knockdown on ribosome biogenesis and translation could impair the regulation of specific gene programs in mouse ESCs. To address this question, control- or Rsl24d1-targeting siRNAs were transfected in CGR8 ESCs followed by RNA-Seq profiling. The top 15% of genes with the most significant expression changes (>1.8 fold change; p < 0.01) were further analyzed. Rsl24d1 loss resulted in the altered expression of 529 genes, including 250 genes upregulated (47%) and 279 genes downregulated (53%) (Table 2). An analysis of Gene Ontology terms associated with genes decreased upon Rsl24d1 depletion revealed a significant enrichment in terms associated to immune response, metabolic processes and transporter activity (p < 0.01) (Fig. 4A, right panel; Supplemental Fig. 4A; Tables 3A and 3B for a full analysis). Conversely, genes upregulated upon si-Rsl24d1 treatments are strongly enriched in terms associated with developmental processes, cell differentiation, cell proliferation and transcription regulation (p < 1E-06) (Fig. 4A, left panel; Supplemental Fig. S4A; Tables 3C and 3D for a full analysis). These results suggest that Rsl24d1 expression in mouse pluripotent ESCs is required to maintain a coordinated regulation of specific transcription programs, including differentiation and development processes, which are most likely important for the control of ESC homeostasis.

**Figure 4.**
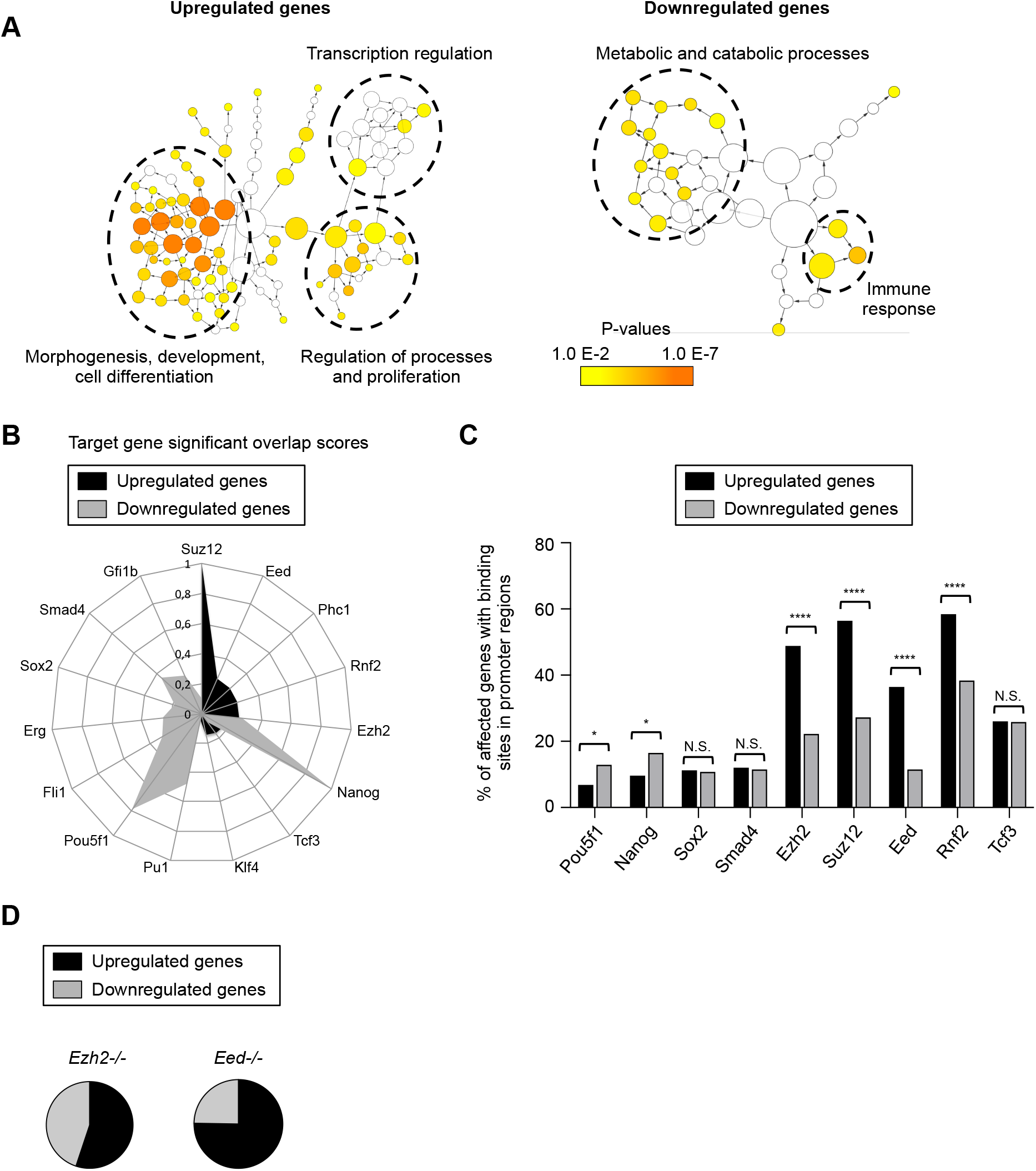
RSL24D1 is required to maintain the regulation of pluripotency and differentiation transcriptional programs. (A) Gene ontology enrichment analysis of biological process terms for genes displaying significant expression changes by RNA-seq analysis in CGR8 cells treated with Rsl24d1 siRNAs. The left and right panels represent hierarchical trees of the most enriched terms in genes up- or downregulated in Rsl24d1-depleted cells, respectively. The size of the nodes represents the numbers of genes associated to each GO term, and the corresponding p-values are indicated by colour codes, according to the scale provided. The most represented GO term categories are indicated. The corresponding data are available in Table S3. (B) Radar plot summarizing the Stemchecker analysis for PTFs and chromatin-associated factors with a significant association score to the 250 upregulated and the 259 downregulated genes in Rsl24d1-depleted ESC^FBS^ CGR8 cells. The corresponding data are available in Table S4. (C) Diagram showing the relative proportion of up- and downregulated genes in Rsl24d1-depleted naive CGR8 cells that contain binding sites in their promoter region (+/-1kb from transcription start sites) for indicated transcription and chromatin-associated factors, established by ChIP-seq analyses (52). P values are indicated (Fisher exact test): * pval<0.05, **** pval<0.0001, N.S.: not significant (pval>0.05). (D) Analysis of the proportion of genes displaying similar expression changes, either up- or downregulation, in Rsl24d1-depleted CGR8 cells and in EED^-/-^ or EZH2^-/-^ ESCs (7-8).

PSCs properties are tightly controlled by distinct epigenetic and transcriptional regulators supporting the expression of programs required for self-renewal and pluripotency while maintaining differentiation programs in a poised state (49, 50). Thus, we hypothesized that the alteration of specific genetic programs upon Rsl24d1 depletion could result from the impaired expression of one or more of these key transcriptional or epigenetic stemness regulators. To test this hypothesis, we first used the StemChecker algorithm to identify ESC-master regulators whose expression is altered upon Rsl24d1 loss (51). Interestingly, this analysis revealed that genes downregulated (n=279) upon Rsl24d1 knockdown were enriched in targets of key PTFs, including Nanog, Pou5f1, Smad4 and Sox2 (Fig. 4B; Table 4A for a full analysis). Conversely, upregulated genes (n=250) were preferentially enriched in targets of essential epigenetic regulators from the polycomb repressive complexes, including Suz12, Eed and Ezh2 from PRC2, and Rnf2 from PRC1 (Fig. 4B; Table 4B for a full analysis).

To further establish whether genes differentially expressed upon Rsl24d1 knockdown are direct targets of these PTFs and PRC factors, we next examined the promoter regions of these 529 genes for enrichment in binding sites for these factors. ChIP-Seq assays performed in murine ESCs, which are available in the ChIP-Atlas database for 9 factors (52), were analyzed to identify binding sites in the vicinity of transcription start sites (TSS ± 1 kb, referred to as promoter regions) for either up- or downregulated genes upon RSL24D1 depletion (Fig. 4C, Table 5). Interestingly, promoters from downregulated genes showed a significant 2-fold enrichment in POU5F1 and NANOG binding sites compared to upregulated genes, therefore supporting the hypothesis that genes downregulated upon RSL24D1 knockdown are enriched in POU5F1- and NANOG-regulated genes. In addition, the promoter regions of upregulated genes were significantly enriched in binding sites for PRC1 and PRC2 factors compared to downregulated genes (Fig. 4C). Strikingly, about half of the promoter regions of upregulated genes were bound by EZH2 (48,9%), SUZ12 (56,4%), EED (36%) or RNF2 (58,4%) in murine ESCs, respectively (Table 5). These results further confirmed that genes down- or upregulated upon RSL24D1 depletion are enriched in targets of POU5F1/NANOG or EZH2/SUZ12/EED/RNF2, respectively.

PRC2 complexes are key epigenetic repressors responsible for the genome-wide deposition of H3K27me2 and H3K27me3 marks to control early embryonic gene expression patterns in ESCs (1, 7-9). Since a large proportion of down- and upregulated genes were associated with PRC2 binding sites, we next analyzed the H3K27 methylation status, established in mouse ESCs (9), of the promoter regions (+/-1kb near TSS) of the 529 genes differentially expressed upon Rsl24d1 depletion (Supplemental Fig. S4B). While H3K27me2 marks were not preferentially enriched near the TSSs of differentially expressed genes, we found that the promoter regions of genes affected by RSL24D1 depletion were less associated with H3K27me1 modifications marking active transcription and were rather significantly enriched in H3K27me3 repressive modifications. These results suggest that the levels of H3K27me3 might be altered when RSL24D1 is depleted. Accordingly, si-Rsl24d1 treated CGR8 cells displayed lower levels of nuclear H3K27me3 compared to si-CTL treated cells (Supplemental Fig. S4C), suggesting that some of the alterations in gene expression detected by RNA-seq upon RSL24D1 knockdown may directly result from this reduction in global H3K27me3.

Finally, we compared RNA-seq predictions from si-Rsl24d1 ESCs with gene expression profiling from EED and EZH2 knockout mouse ESCs, respectively (7, 8). Interestingly, genes displaying an altered expression upon RSL24D1 depletion were enriched in genes that are mis-regulated in EED^-/-^ or EZH2^-/-^ ESCs (Supplemental Fig. 4E). Strikingly, genes with an expression profile that was similarly impaired in PRC2 mutant cells and RSL24D1-depleted cells were predominantly upregulated or derepressed genes (Fig. 4E). Altogether these results suggest that genes up- or downregulated upon RSL24D1 knockdown are likely controlled by distinct molecular mechanisms. On the one hand, downregulated genes are enriched in key PTF target genes, suggesting that the transcriptional regulatory activities of POU5F1 and NANOG are decreased in si-Rsl24d1 treated ESCs. On the other hand, the enrichment of H3K27me3 sites and binding sites for PRC2 proteins in promoter regions of upregulated genes suggests that RSL24D1 depletion most likely hinders the repressive activity of PRC2. This results in lower H3K27me3 deposition, therefore leading to the premature activation of developmental genes normally silent in pluripotent ESCs.

### The translational regulation of key stemness factors is impaired upon RSL24D1 depletion in mouse ESCs

In order to investigate how RSL24D1 depletion could affect the activity of PTFs and PRC2 factors, we first measured whether the loss of RSL24D1 affected their transcription. Interestingly, the expression of these PTFs and PRC factors was not strongly impaired at the RNA level in si-Rsl24d1 ESCs, except for Nanog (Supplemental Fig. S5A), suggesting that rather the production and/or the activity of the corresponding proteins might be affected upon RSL24D1 depletion.

Since RSL24D1 depletion decreases global protein synthesis, we hypothesized that the alteration of PRC2 activity could result from a perturbation of Eed, Ezh2 and Suz12 mRNA translation. To address this question, mRNAs associated with the different ribosomal fractions, and in particular with actively translating ribosomes (i. e. polysomes), were analyzed by RT-qPCR assays in CGR8 cells treated with either si-CTL or si-Rsl24d1 (Supplemental Fig. S5B and S5C). As previously described, downregulating RSL24D1 in ESCs impaired the accumulation of the cytoplasmic 40S, 60S and 80S ribosome fractions (Supplemental Fig. S5B). As expected, the majority (>92%) of Eed, Ezh2 and Suz12 mRNAs detected in this assay were associated with ribosome fractions corresponding to polysomes suggesting that these mRNAs were actively translated in self-renewing ESCs (Fig. 5A and Supplemental Fig. S5C, gray curves). However, in RSL24D1-depleted cells, the association of Eed and Ezh2 mRNAs with polysomal fractions was significantly reduced and correlated to an increased detection in fractions corresponding to free mRNPs and monosomes (Fig. 5A and Supplemental Fig. S5C, black bar graphs). In contrast, the polysome versus free mRNPs/monosome comparison was not statistically significant for Suz12 mRNAs (Fig. 5A), in particular due to a biological variability in the detection of these mRNAs throughout gradients. However, analyses of the polysome fractions alone seemed to indicate that Suz12 mRNAs also tend to transit from heavy to light polysomes upon RSL24D1 depletion (Supplemental Fig. S5C).

**Figure 5.**
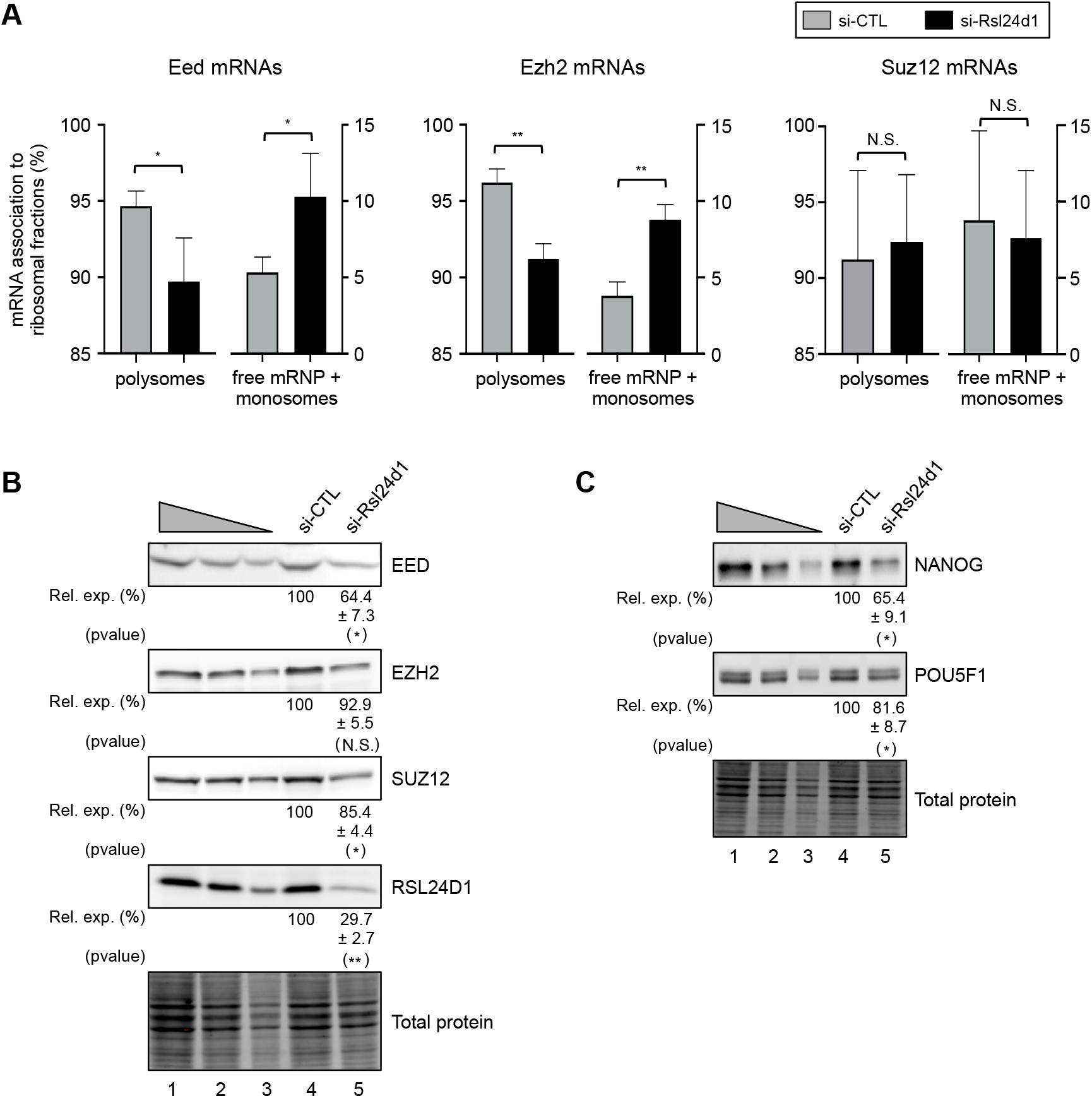
RSL24D1 depletion impairs the translation of core PRC2 factors and the expression of key PTFs. (A) Relative proportions of Eed, Ezh2 and Suz12 mRNAs detected by qRT-PCR in free mRNP, monosomal or polysomal fractions, based on quantifications detailed in the supplemental Figure S5C, in CGR8 ESC^FBS^ treated with CTL-(grey) or Rsl24d1-targeting siRNAs (black) (n=3). P values are indicated (paired two-tailed Student’s t-test): ** pval<0.005, * pval<0.05, N.S.: not significant (pval>0.05). (B) Representative immunoblots of RSL24D1 (n=3) and 3 PCR2 proteins (EED n=3, EZH2 n=3, SUZ12 n=4) in CGR8 ESC^FBS^ treated with non-targeting or Rsl24d1-targeting siRNAs. Lanes 1 to 3 correspond to serial dilutions of si-CTL-treated ESC^FBS^ (1:1, 1:3 and 1:9, respectively). TCE labeling of tryptophan-containing proteins (referred as Total Proteins) is used for normalization. Quantifications of RSL24D1, EED, EZH2 and SUZ12 signals normalized to total proteins and relative to the si-CTL-treated conditions (Rel. exp. (%)) are indicated below each panel. P values are indicated and relative to the si-CTL-treated conditions (paired two-tailed Student’s t-test): ** pval<0.01, * pval<0.05, N.S.: not significant (pval>0.05). (C) As previously described for panel B for POU5F1 (n=3) and NANOG (n=3) in CGR8 cells treated with non-targeting (si-CTL) or Rsl24d1-targeting siRNAs. * pval<0.05.

To further confirm that RSL24D1 downregulation impaired the translation of Suz12, Eed and Ezh2 mRNAs, steady-state expression levels of the corresponding proteins were assessed by semi-quantitative western blots from CGR8 total cell extracts. Interestingly, RSL24D1 downregulation induced a significant reduction in the accumulation of EED and SUZ12 proteins, albeit no significant decrease in expression was observed for EZH2 (Fig. 5B). Altogether, these observations suggest that the loss of PRC2 activity upon RSL24D1 depletion, which may cause both a defect in H3K27me3 and an upregulation of specific gene programs associated with development and differentiation, is likely caused by a defect in the translation of Eed, Ezh2 and Suz12 mRNAs.

Similar to PRC2 factors, we hypothesized that a translational decrease of POU5F1 and NANOG mRNAs may be responsible for the downregulation of specific genetic programs enriched in POU5F1 and NANOG target genes. This hypothesis was indeed confirmed by semi-quantitative western blots that demonstrated a significant reduction in the detection of both POU5F1 and NANOG proteins in si-Rsl24d1 treated cells compared to si-CTL cells (Fig. 5C). Immunostaining experiments confirmed a global and significant loss of POU5F1 expression in si-Rsl24d1-treated ESCs suggesting that the POU5F1 downregulation previously observed by western blot in total cell extracts did not result from the emergence of POU5F1 negative cells in ESC colonies but rather resulted from a global decrease in POU5F1 expression (Supplemental Fig. S5D). Altogether, these results confirm that RSL24D1 depletion directly impairs the translation and the accumulation of both key PTFs and core PRC2 factors, which respectively play pivotal roles in controlling gene expression programs and the chromatin landscape underlying cell fate decisions in pluripotent ESCs.

### High RSL24D1 expression supports the maintenance of self-renewal capacities but is dispensable for ESC differentiation

Since transient RSL24D1 depletion altered the expression of several PTF target genes and promoted the activation of genes involved in developmental programs, we examined the functions of RSL24D1 for ESC fundamental self-renewal and differentiation capacities. First, stable RSL24D1 downregulation by constitutively expressed shRNAs significantly impaired, in a dose-dependent manner, the proliferation of CGR8 cells in both real-time assays (Fig. 6A) and long-term kinetics of proliferation (Fig. 6B). Accordingly, expression of sh-Rsl24d1-2 demonstrating the most significant loss of RSL24D1 (Supplemental Fig. S3B and S6A) also correlated with the strongest proliferation impact on CGR8 cells (Fig. 6A and 6B). Considering that ESC self-renewal capacities are tightly controlled at the molecular level by key transcriptional programs, we next analyzed the expression of several major PTFs upon stable depletion of RSL24D1 in CGR8 cells. In contrast to si-Rsl24d1 transient knockdowns, RT-qPCR assays revealed a significant downregulation, at the mRNA level, of Nanog, Pou5f1 and Sox2 in cells expressing the sh-Rsl24d1-2 sequence while Klf4 expression remained unaffected (Supplemental Fig. S6A). The expression of the shRsl24d1-1 sequence induced a lower depletion of RSL24D1 correlated to less pronounced alterations of these PTFs, with only Sox2 being downregulated and Klf4 slightly upregulated (Supplemental Fig. S6A). Using western blot assays, a significant downregulation of POU5F1 was also confirmed in sh-Rsl24d1-2 expressing CGR8 cells (Supplemental Fig. S6B), suggesting that RSL24D1 stable depletion likely affected ESC self-renewal capacities. To determine whether this is the case, CGR8 cells expressing non-targeting shRNAs (sh-Control), or shRNAs specifically targeting Pou5f1 or Rsl24d1 mRNAs were seeded at clonal density to recapitulate the formation of individual undifferentiated colonies displaying high levels of alkaline phosphate (AP) activity (53). As expected from its key role in controlling ESC self-renewal transcriptional programs, POU5F1 downregulation led to a drastic reduction (>90 %) in the number of AP positive colonies compared to control shRNA treated cells (Fig. 6C and Supplemental Fig. S6C). Similarly, RSL24D1 depletion caused a dose-dependent and significant reduction (>50%) of AP-positive ESC colonies (Fig. 6C and Supplemental Fig. S6C), suggesting that a high expression of RSL24D1 in mouse ESCs is required to support both self-renewal and proliferation.

**Figure 6.**
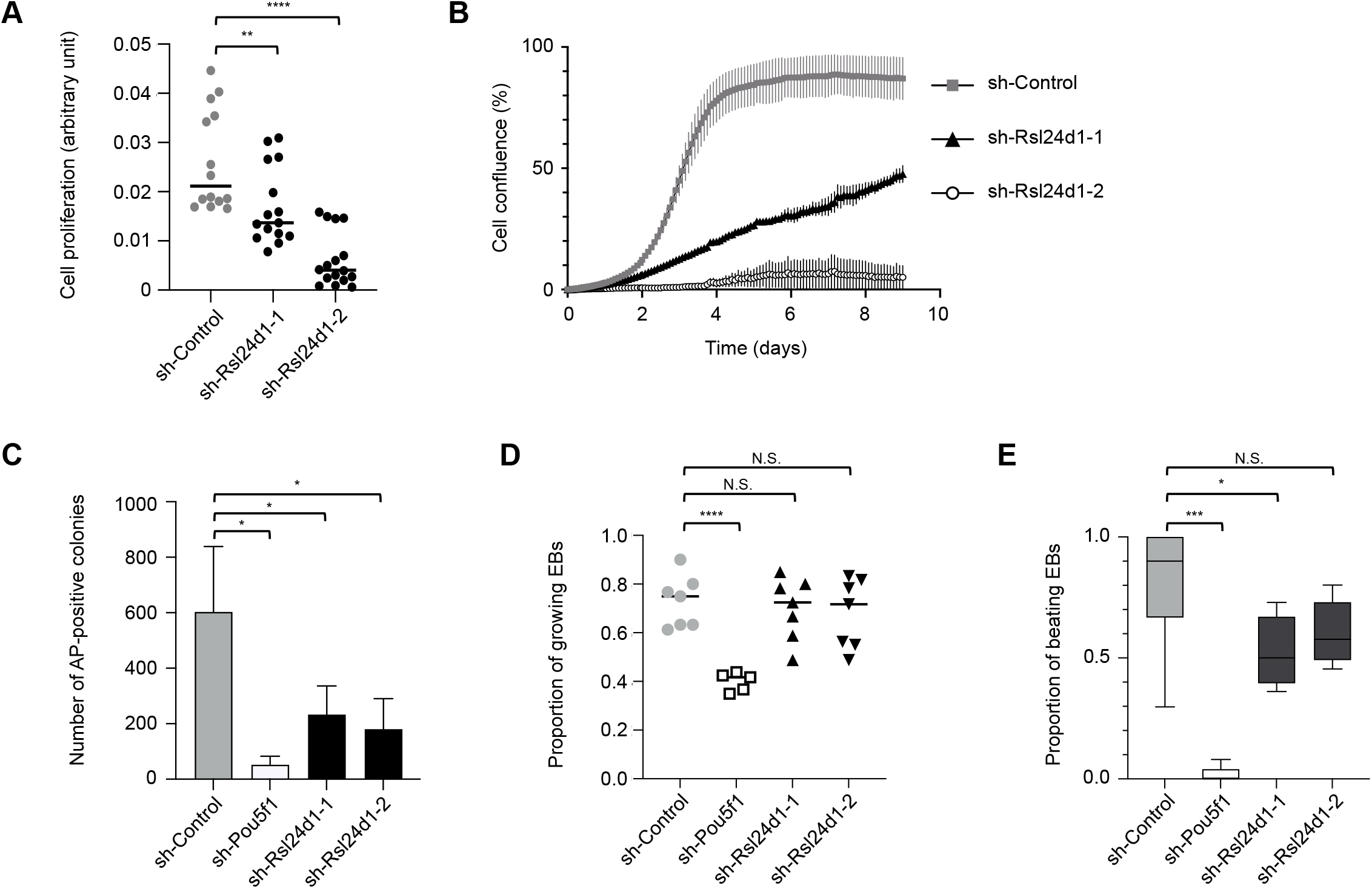
RSL24D1 is required for mouse ESC proliferation and self-renewal. (A) Analysis of proliferative capacities defined by the cell growth index quantified by the xCELLigence RTCA technology in the first 12 hours after plating CGR8 cells in ESC^FBS^ conditions. Cells were infected with lentiviral vectors expressing a control non-targeting shRNA (grey dots, sh-Control, n=14 wells), or 2 distinct shRNAs targeting Rsl24d1 (black dots, sh-Rsl24d1-1, n=15 wells; sh-Rsl24d1-2, n=16 wells). P values are indicated and relative to the sh-control condition (unpaired two-tailed Student’s t test): ** pval<0.005, **** pval<0.0001. (B) Long term proliferation estimated by cell confluency analysis of naive CGR8 cells expressing a non-targeting shRNA (sh-Control, grey squares) or 2 distinct shRNAs targeting Rsl24d1 (sh-Rsl24d1-1, black triangles and sh-Rsl24d1-2, white circles)(n=6). (C) Analysis of the self-renewing capacities of CGR8 cells in ESC^FBS^ conditions estimated by the quantification of individual colonies with alkaline phosphatase activity detected by colorimetric labelling (corresponding raw data are detailed in Supplemental Fig. 6C). CGR8 colonies expressing control-(n=4 wells), Pou5f1-(n=4 wells) or Rsl24d1-targeting shRNAs (sh-Rsl24d1-1, sh-Rsl24d1-2) (n=5 wells) were compared. * pval<0.05, unpaired two-tailed Student’s t test. (D) Analysis of *in vitro* differentiation capacities of CGR8 cells expressing control-(n=7 wells), Pou5f1-(n=5 wells) or Rsl24d1-shRNAs (n=7 wells) defined by the proportion of seeded embryoid bodies surviving and growing during a 12-day differentiation course. **** pval<0.0001, N.S.: not significant (pval>0.05), unpaired two-tailed Student’s t test. (E) Analysis of the proportion of 12-day old EBs, generated from CGR8 cells expressing control-(n=7 wells), Pou5f1-(n=5 wells) or Rsl24d1-shRNAs (n=7 wells), containing functional self-beating cardiomyocytes. *** pval<0.001, * pval<0.05, N.S.: not significant (pval>0.05), unpaired two-tailed Student’s t test.

Finally, we compared the differentiation capacities of sh-Control, sh-Pou5f1 and sh-Rsl24d1-1/2 expressing CGR8 cells by EB differentiation assays. As expected, POU5F1 knockdown strongly impaired the capacities of CGR8 cells to form viable 10 day-old EBs, whereas RSL24D1 downregulation had no significant impact on EB formation compared to sh-Control ESCs (Fig. 6D). We next assessed the functional differentiation of EB-forming cells by counting the proportion of EBs displaying spontaneous beatings, which characterize the presence of fully differentiated cardiomyocytes (54). This analysis revealed that POU5F1 depletion almost completely abolished EB beating (Fig. 6E), likely reflecting its role in early cell fate determination (55-57). Conversely, RSL24D1 stable knockdowns only partially impaired the formation of functional cardiomyocytes after 10 days of differentiation (Fig. 6E). Thus, the depletion of RSL24D1 affects ESC self-renewal and proliferation capacities but does not seem however to significantly interfere with the processing of differentiation.

## Discussion

Despite displaying reduced translation activity compared to differentiated progenies (16, 17, 19), ESCs have also been previously shown to paradoxically express RBFs and RPs at higher levels than differentiated cells (28, 30, 31, 58). These observations therefore suggest that naive ESCs may need to accumulate, at a steady-state level, a pool of ribosomes sufficient to sustain the rapid proteome changes and increased translational activities required to achieve all possible programs of differentiation in response to environmental signals. In order to better define the molecular mechanisms and factors coordinating the highly regulated ribosome production in ESCs, we first identified RBFs enriched in naive ESCs compared to differentiated cells. Our study revealed that RSL24D1, a homolog of the yeast Rlp24 LSU ribosome biogenesis factor, is consistently expressed at high levels in PSCs, including mouse ESCs and iPSCs, and is rapidly downregulated after differentiation induction.

Since RSL24D1 function was not established in higher eukaryotes, we next determined that RSL24D1 is predominantly localized in nucleoli, which are nuclear domains playing a central role in the transcription and maturation of pre-rRNAs (59). Considering the strong conservation of RSL24D1 structure and functions relative to its yeast homolog Rlp24, it is also likely that RSL24D1 shuttles between the nucleus and the cytoplasm (42). Accordingly, immunostaining assays revealed that, while the majority of endogenous RSL24D1 is detected in the nucleoli of mouse ESCs (Fig. 2B), RSL24D1 signals are also detected in the cytoplasm, in contrast with FBL signals strictly detected in the nucleoli. Conversely, RPL24 is almost exclusively present in the cytoplasm as described in yeast (42). In yeast, Rlp24 is assembled at the initial steps of pre-60S maturation corresponding to the state B (60), together with Tif6, Nog1 and Mak11 (23, 61), and then allows the recruitment of the hexameric Drg1 AAA-ATPase (42, 62). As the particles exit the nucleus, Drg1 gets activated by nucleoporins and releases Rlp24 from the pre-60S by mechanical force (42, 62-64), therefore allowing its substitution by the canonical ribosomal protein Rpl24. Although the loss of Rlp24 in yeast induced moderate alterations of the 35S and 27SB rRNA intermediates, we did not observe any significant modifications of the rRNA processing in mouse ESCs treated with siRNAs targeting Rsl24d1 mRNAs (data not shown). Even though this observation may reflect a difference in the efficacy of RSL24D1/Rlp24 depletions in each model, both the conditional Rlp24 mutant in yeast (42) and the depletion of RSL24D1 in human cells (65) did not strongly impact the accumulation of maturation intermediates or mature LSU rRNAs. These observations indeed suggest that RSL24D1 rather guides the association of other proteins to early states of pre-60S, in particular Tif6, the yeast homolog of EIF6, rather than regulating early exonucleolytic and endonucleolytic rRNA processing (42, 60, 66). Consistently, the depletion of RSL24D1 in mouse ESCs is correlated to a loss of EIF6 association to nuclear pre-ribosomes (Fig. 3D). Interestingly, RSL24D1 and EIF6 show different dynamics of disassembly from cytoplasmic ribosomes, consistent with their dynamics of assembly/release in yeast (Fig. 2C). EIF6 dissociation seems to occur belatedly as it is more stably associated with cytoplasmic ribosomal particles. These observations are therefore consistent with the yeast model describing Tif6 removal as the latest maturation step of 60S particles (23, 67). Furthermore, yeast Rlp24 has a key protein-protein mimicry role during pre-60S maturation until it is replaced, after nuclear export of pre-LSU particles, by the canonical Rpl24, which is strictly localized to the cytoplasm (42). Our data strongly support a conserved mimicry function for RSL24D1 in mouse ESCs as its depletion impairs the association of RPL24 with cytoplasmic ribosomes (Fig. 3D).

Moreover, consistent with a conserved role in the LSU biogenesis, RSL24D1 depletion in mouse ESCs caused a loss of 60S and 80S subunits relative to 40S particles. A similar imbalance of the accumulation of 60S and 80S over 40S subunits was previously observed in yeast upon Rlp24 depletion (42) or 60S biogenesis repression (33, 68). In addition, we detected the presence of half-mers for di- and tri-ribosomes in si-Rsl24d1 treated cells (Figure 3B), which were previously observed upon LSU biogenesis alterations in yeast (69, 70). Finally, RSL24D1 has also been detected in nuclear pre-60S cryo-structures while it is absent from cytoplasmic pre-LSU in human cells and we demonstrated that RSL24D1 expression profile is also conserved in human ESCs, further supporting a conserved molecular function in ribosome biogenesis and stem cells throughout evolution.

We then established that RSL24D1 depletion, in addition to impairing the accumulation of 60S subunits in the cytoplasm, also strongly alters global translation in naive CGR8 cells. This result agrees with previous observations that the loss of expression of either HTATSF1, which controls rRNA processing, or SSU biogenesis factors also impair translation in mouse ESCs (21, 31). In addition to a global translational impact, we provided evidence that RSL24D1 depletion is correlated to a reduced expression of specific instable proteins including POU5F1 and NANOG. It is worth noting that the decrease in NANOG expression may result from a combination of both transcriptional and translation defects while POU5F1 downregulation was not detected at the transcriptional level in the same conditions (Fig. 5C and Supplemental Fig. 5A). A highlight of this study is that the steady-state level of several PRC2 factors, including EZH2, EED and SUZ12 is tightly controlled by the level of ribosome biogenesis in ESCs. Although these factors play a key role in organizing the landscape of active and transcriptionally silent chromatin, and are essential to repress developmental genes by maintaining H3K27me3 modifications on promoter regions, their activity rather seems rate-limiting as slight changes in expression have a massive impact on global gene regulation in ESCs. Indeed, the downregulation of RSL24D1 in mouse ESCs induced a reduction in Eed, Suz12 and Ezh2 mRNAs association to polysomes, which is predicted to have a direct impact on corresponding neosynthesized proteins. Accordingly, despite RSL24D1 depletion causing a reduction ranging from 7% to 35% in corresponding EED, SUZ12 and EZH2 protein levels, we observed a correlated and significant increased expression, at the transcriptional levels, of hundreds of PRC2 target genes normally associated to repressive H3K27me3 marks. Many of these PRC2 target genes correspond to developmental genes, including members of the HOX (HOXB1, 2, 3, 9 and C13) and WNT families (WNT2B, 3, 10A and 10B), transcription factors such as NKX1-2, NKX2-9, POU4F3, SALL3 and NEUROD1. Altogether, this therefore suggests that the versatile status of ESCs, which must rapidly switch from a transcriptionally active self-renewal program to the activation of poised developmental or differentiation programs, requires a finely tuned translation of PRC2 factors that is coordinated with ribosome biogenesis in ESCs.

At the physiological level, we showed that RSL24D1 depletion caused a significant loss of proliferation and self-renewal capacity while having a moderate impact on ESC differentiation *in vitro*. This is in agreement with RSL24D1 expression profile, which is high in ground state and naive pluripotent ESCs but low in differentiating progenies, therefore supporting the hypothesis that RSL24D1 expression is required for ESC maintenance. Surprisingly, attempts to overexpress an exogenous FLAG-RSL24D1 protein at near ESC endogenous levels, induced a rapid and stable downregulation of the endogenous RSL24D1 in ESCs (data not shown). This therefore suggests that PSCs finely tune the steady-state level of RSL24D1, most likely to avoid aberrant RSL24D1 levels that could be detrimental for ribosome biogenesis and stem cell homeostasis.

Altogether, we propose a model where RSL24D1 actively contributes to the biogenesis of pre-60S ribosomal particles and to the accumulation of mature monosomes and polysomes in the cytoplasm of naive mouse ESCs (Fig. 7). We suggest that RSL24D1 shuttles back and forth between the nucleoli and the cytoplasm to support an elevated rate of 60S biogenesis in ESCs while its activity is less required in differentiating ESC progenies. Here, we demonstrated for the first time that the expression of a RBF of the LSU is regulated as ESCs transition from naive to differentiated states, and is required to maintain ESC self-renewal and proliferation. Interestingly, RSL24D1-mediated ribosome biogenesis therefore appears to have a dual role in controlling ESC self-renewal. On the one hand, RSL24D1 maintains a balanced expression of key PTFs, including POU5F1 and NANOG, to control pluripotency transcriptional programs. Furthermore, RSL24D1 expression profile is correlated to POU5F1 expression during CGR8 differentiation (Supplemental Fig. 1C), and RSL24D1 is a predicted target gene of POU5F1 (Table S4 and ChIP-Atlas database) and NANOG (ChIP-Atlas database). The stable depletion of POU5F1 by shRNAs is correlated to a significant reduction (>60%, data not shown) in RSL24D1 expression at the mRNA and protein levels in CGR8, suggesting that RSL24D1 expression could be directly controlled by POU5F1 or NANOG in mouse ESCs. On the other hand, RSL24D1 sustains steady-state translation of PRC2 factors and therefore maintains repressive H3K27me3 chromatin marks to prevent the premature activation of developmental genes in naive ESCs (Fig. 7). Intriguingly, EZH2 was recently shown to have a PRC2-independent function by promoting the interaction between NOP56 and FBL in human cancer cells and mouse extraembryonic endoderm stem cells, and enhancing 2’-O-methylation of rRNAs (71). This observation raises the question of whether RSL24D1 might influence rRNA 2’-O-methylation by differentially modulating EZH2 expression in naive and differentiated ESCs. Altogether, these results therefore further support that RSL24D1 may be at the core of a regulatory loop that engages the regulation of ribosome biogenesis and translation with chromatin epigenetic and transcription regulations in order to precisely control ESC self-renewal and pluripotency properties.

**Figure 7.**
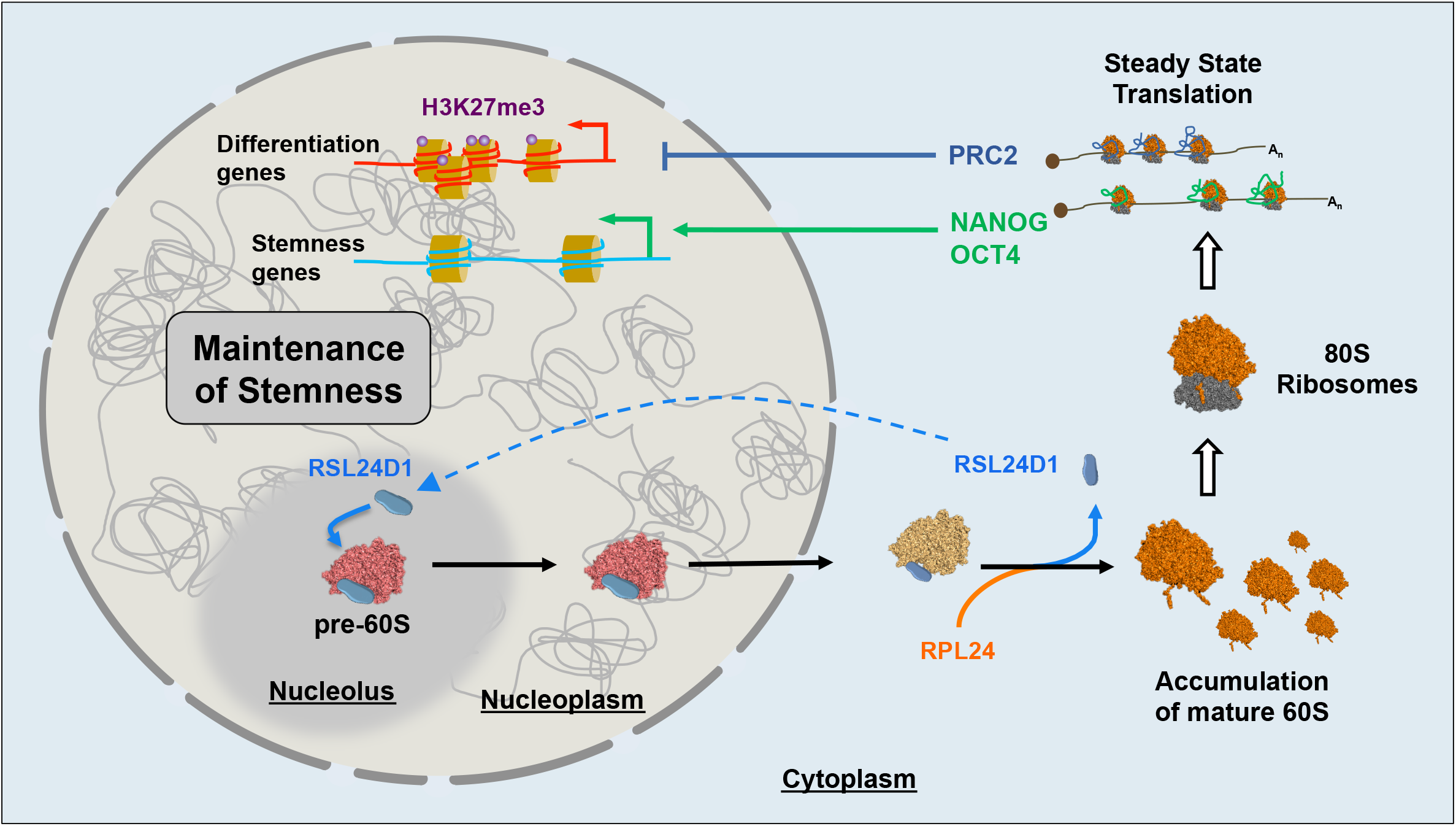
**Model describing the functions of RSL24D1 in the LSU biogenesis and in the translational regulation of key pluripotency and developmental programs in mouse ESCs.**

Finally, modulating the production of ribosomes might not only impact global translation but could also be a mechanism to control the translation of specific subsets of mRNAs that regulate ESC fate. Indeed, it is puzzling that despite an active ribosome biogenesis, ESCs display a globally reduced translational activity, which is correlated to a relatively small cytoplasm compared to differentiated cells. A first hypothesis could be that mouse ESCs have a short cell cycle and therefore an elevated cell division rate that could stimulate ribosome biogenesis. However, whether all the ribosomes present in PSCs are actively engaged in translation still remains to be established. Another seducing hypothesis to explain this discrepancy may be that the ribosome concentration may regulate substrate selectivity and that maintaining a high ribosome biogenesis, and therefore concentration allows the translation of specific programs. This “Ribosome Concentration” model has been previously discussed (72) and may explain why the translation of ESC-relevant mRNAs requires such an elevated amount of ribosomes. One could therefore speculate that fluctuations of ribosome biogenesis, i.e. upon RSL24D1 expression decrease, could affect specific translation programs. Consistently, several observations indicate that upstream open reading frames (uORFs) are commonly used in naive ESCs compared to differentiated cells (73), and that the mRNAs coding for key PTFs, including Nanog, possess multiple uORFs which specifically enhance their translation in ESCs (74). Hence, uORF-carrying mRNAs may require a certain amount of ribosomes to be correctly translated in the context of ESCs while other non-uORF mRNA may be repressed in this context. Transposed to the ESC model which naturally modulates the production of ribosomes depending on their cellular fate, one could propose that naive and differentiating ESCs differentially modulate the translation of specific populations of mRNAs in addition to the global regulation of translation, as a mean to control specific gene programs, such as Nanog-dependent pluripotency or PRC2-dependent differentiation programs. Along these lines, the direct impact of ribosomes on gene-specific regulations has been highlighted in the context of congenital diseases, the ribosomopathies, which are caused by mutations in RBFs or RPs, and characterized by quite heterogeneous phenotypes ranging from specific developmental alterations to increased risks of cancer (72). Therefore, better defining the molecular feedbacks between ribosome biogenesis, translation and additional key steps of gene expression, including chromatin modifications and transcription, could not only benefit to a developmental research but also to open novel avenues to consider disease treatments.

## Materials and methods

### RNA-seq datasets

RNA-seq data for mouse pluripotent cells, differentiated cells and tissues were obtained from a previously published dataset (GSE45505) (36).

### Cell culture

CGR8 mouse embryonic stem cells (ECACC General Collection, 07032901) were cultured on 0.2% gelatin-coated plates either in ESC^FBS^ conditions using GMEM BHK-21 (Gibco) supplemented with 10% ESC-grade heat-inactivated Fetal Bovine Serum (FBS), 1x non-essential amino acid (Gibco), 2 mM Sodium Pyruvate (Gibco), 100μM β-mercaptoethanol (BME, Sigma-Aldrich), 10^3^U/ml Leukemia Inhibitory Factor (LIF, StemCells Technologies) or in ESC^2i^ conditions using 45% DMEM/F12 (Gibco), 45% Neurobasal medium (Gibco), 100μM BME, 1.65% Bovine Serum Albumin Fraction V (Gibco), 1X penicillin-streptomycin (Gibco), 1% N2 supplement (Gibco), 2% B27 supplement (Gibco), 1X ESGRO 2i supplement (MEK1/2, GSK3β inhibitors, Millipore) and 10^3^U/ml Leukemia Inhibitory Factor (Millipore).

The mouse ESC lines R1 (ATCC-SCRC-1011) and G4 (RRID:CVCL_E222) were cultured on irradiated mouse embryonic fibroblasts, in High Glucose DMEM (Gibco), 10% ESC-grade heat-inactivated FBS, 1X non-essential amino acid, 1mM Sodium Pyruvate, 100μM BME and 10^3^U/ml LIF.

H9 and OSCAR human embryonic stem cells were cultured as previously described (41) and were provided by the Savatier laboratory from the Stem cell and Brain Research Institute (Inserm U1028), 69500 Bron, France.

HEK-293T cells were cultured in DMEM (Invitrogen) supplemented with 10% heat inactivated FBS (Sigma-Aldrich), 1X non-essential amino acid, 2mM Sodium Pyruvate and 1X penicillin-streptomycin.

All cell lines used in the study were confirmed mycoplasma-free (Lonza, MycoAlert kit).

### MEF reprogramming into iPSCs

MEFs were isolated from R26^rtTA^;Col1a1^4F2A^ E13.5 embryos after removal of the head and internal organs (75). The remaining tissues were physically dissociated and incubated in trypsin 10 minutes at 37°C. Dissociated cells were resuspended in MEF medium (DMEM supplemented with 10% fetal bovine serum, 100 U/mL penicillin / streptomycin, 1mM sodium pyruvate, 2mM L-glutamine, 0.1mM Non Essential Amino Acids and 0.1mM β-mercaptoethanol).

To induce the reprogramming process, MEFs were plated in six-well plates at 80,000-100,000 cells per well in MEF medium (75). The following day the medium was replaced by fresh MEF medium containing 2μg/mL doxycyclin. MEFs were reseeded after 72h on 0.1% gelatin-coated plates in iPSC medium (DMEM containing 15% KnockOut Serum Replacement, 1,000 U/mL leukemia inhibitory factor, 100 U/mL penicillin / streptomycin, 1mM sodium pyruvate, 2mM L-glutamine, 0.1mM Non Essential Amino Acids and 0.1mM β-mercaptoethanol). Every day, medium was either replaced by or supplemented with doxycyclin-containing fresh medium.

After 14 days of reprogramming, iPS colonies are picked individually, amplified and maintained for several passages.

### Differentiation assays

Embryoid body (EB) formation assays were performed by the hanging drop method (76, 77). Briefly 400 cells were cultured in 20 μL hanging drops for 2 days in GMEM BHK-21 supplemented with 20% heat-inactivated FBS, 1x non-essential amino acid, 2mM Sodium Pyruvate, 100μM BME. EBs were then collected in non-adherent culture dishes and cultured for 3 additional days. At Day 5, cell aggregates were cultured on 0.2% gelatin-coated dishes in the presence of 10nM retinoic acid (Sigma-Aldrich) and the media was changed every 2 days.

### Plasmids

pLKO.1-puro plasmids (Addgene #8453) were cloned as previously described (78). Briefly, the pLKO.1 plasmid was digested with EcoRI and AgeI restriction enzymes. Linear plasmids were ligated with annealed oligomers containing the shRNA sequence flanked by EcoRI and AgeI restriction sites, respectively. All clones were sequenced to confirm the insertion of each shRNA sequence in the pLKO.1 backbone prior to lentiviral production.

### Lentiviral production and infection

The production of lentiviral particles for shRNA expression was performed as previously described (79, 80). Briefly, HEK-293T cells were co-transfected with corresponding pLKO.1 plamids and 3^rd^ generation packaging lentiviral plasmids (pLP1, pLP2, pLP-VSVG) using FuGENE HD (Promega), according to the manufacturer’s recommendations. Cell media containing lentiviral particles were concentrated using centricon column (Vivaspin 20, Sartorius). Each viral production was titrated beforehand to establish a multiplicity of infection (MOI) sufficient to provide a complete resistance to 1.5 µg/mL of puromycin (Sigma-Aldrich).

ESCs were infected with an MOI=1 for 16 hours in ESC^FBS^ conditions supplemented with polybrene at 8 µg/mL (Sigma-Aldrich). 24h after infection, cells were selected with 1.5 µg/mL of puromycin for at least 48 hours. shRNA sequences used in this study are reported in Table S6A.

### siRNA Transfection

ESCs were transfected with 20nM siRNAs using DharmaFECT 1 transfection reagent (Horizon Discovery) according to the manufacturer’s protocol, for the indicated times in ESC^FBS^ conditions. Negative control pool (D-001810-10-20, ON-TARGETplus, Horizon Discovery) was used as a negative control for siRNA transfection while RSL24D1 knockdowns were achieved using ON-TARGETplus RSL24D1 SMARTpool (L-054445-01-0010, Horizon Discovery).

### Colony-formation assay

ESCs were plated at clonal density (60 to 100 cells per cm2) on gelatin-coated plates in ESC^FBS^ conditions. The medium was changed every two days for seven days before detection of alkaline phosphatase-positive colonies (Alkaline Phosphatase detection kit, Merck-Millipore). Alkaline phosphatase-positive colonies were analyzed and quantified using ImageJ analysis software (81).

### Proliferation assays

The proliferation was first assessed using the x-CELLigence Real-Time Cell Analysis (RTCA) system (ACEA Biosciences, San Diego, CA, USA). 5000 cells were seeded in a 96 E-plates (Roche) and the electrical impedance was acquired every 15 minutes for seven days to establish a cell index value extracted from the linear regression of the proliferation curve. The data were analyzed by RTCA software.

In addition, the proliferation was also monitored using the high-definition automated imaging system IncuCyte (Essen BioScience), according to the manufacturer’s instructions. 750 infected and selected cells were seeded in a 24-well plate, with 2-hour interval snapshots. Proliferation rates were estimated according to the manufacturer’s insctructions.

### Western Blots

Cells were lysed in 1x Laemli buffer (50mM Tris-HCl pH 6.8, 2% SDS, 5% BME, 10% glycerol and 0.05% bromophenol blue) and analyzed by SDS-polyacrylamide gel electrophoresis and western blotting using antibodies indicated in Table S7B.

For total protein quantification, 0.5% trichloroethanol (Sigma-Aldrich) was included in the SDS-polyacrylamide gel prior to electrophoresis and activated post-electrophoresis with UV light for 45s. Chemiluminescent and fluorescent signals were acquired on ChemiTouch MP imaging system (Bio-rad) and quantified using ImageLab software (Bio-rad) by normalizing signals of interest to housekeeping protein (β-actin, GAPDH) or total protein signals. Serial sample dilutions were systematically loaded onto gels and analyzed to verify the linearity of quantified signals.

### Immunofluorescence assays and High-Content analysis System (HCS)

ESCs were cultured in ESC^FBS^ conditions either on 0.2% gelatin-coated coverslip (SPL) or gelatin-coated 96-well plates. 48 after siRNA transfections, ESCs were fixed for 10 min in 4% formaldehyde and permeabilized for 10 min in 0.1% Triton X-100. Cells were then incubated 1 hour in blocking solution (1X PBS, 0.1% tween 20, 5% BSA) and then incubated with primary and secondary antibodies indicated in Table 6B. Images were acquired using a Zeiss Axio Imager M2 microscope coupled with the Zen 2 Pro software (Zeiss) and processed with ImageJ.

For deeper statistical results, cells were plated on a 96 well carrier plate (Perkin Elmer) optimized for sensitive and resolved fluorescence microscopy. At least 90% of the well surface is acquired in one non-confocal plane with Harmony software on an Operetta CLS Flex High-content-Screening system (Perkin Elmer) equipped with 20x/NA1.0 water objective. The set-up was optimized to reach at least a difference of 10000 fluorescence levels between the noise and the signal of interest to allow a robust images analysis and quantification. Data were analyzed with Columbus software (Perkin Elmer). Briefly, on the image of full wells, colonies were located on the nuclei labeling with the appropriate tuned find image region algorithm. To discard small and large colonies, areas in the range of 500 to 40 000 µm^2^ of cells were selected. Then cells in each colony were found with the appropriate tuned find nuclei algorithm. Cell debris and objects with more than one nucleus were excluded by filtering the nuclei according to roundness and surface. Fluorescence intensities are calculated for each selected cellular region for all wells.

### Cell fractionation

ESCs were collected by trypsinisation and gently lysed for 10 min on ice in hypotonic buffer (HB) containing 10 mM KCl, 0.5 mM MgCl_2_, 10 mM Tris-HCL pH 7.4, 1X cOmplete EDTA-free protease inhibitors ™ (Roche) and 1U/µL RNAseOUT (Invitrogen). 0.02% NP-40 was subsequently added for 5 more minutes and nuclei and cytoplasm were then separated through sequential centrifugations. Nuclei were washed with HB supplemented with 0.01% NP-40 and resuspended in Buffer A (250 mM Saccharose, 250 mM KCl, 5mM MgCl_2_ and 50mM Tris-HCl pH7.4) with DNAse I (2000U/mL). Cytoplasmic fractions in HB were adjusted to 250 mM KCL.

### Ribosome purification on sucrose cushion

Cytoplasmic and nuclear fractions were loaded on 1 mL sucrose cushion (1M saccharose, 250mM KCl, 5mM MgCl_2_ and 50mM Tris HCl pH7.4) and centrifugated at 250.000 g for 2 hours at 4°C. Pellets were washed twice with cold water and resuspended in Buffer C (25mM KCl, 5mM MgCl_2_ and 50 mM Tris-HCl pH 7.4).

### Polysome profiling

ESCs were treated with 25 µg/mL of emetine (Sigma-Aldrich) for 15 min and lysed in 10 mM Tris-HCL pH7.5, 5 mM MgCl_2_, 100 mM KCl, 1% Triton X-100, 2 mM DTT, 1U/μL RNAseOUT and 2X cOmplete EDTA-Free protease inhibitors. Lysates were centrifugated at 1300 g for 10 min to pellet nuclei. Supernatants corresponding to cytoplasmic fractions were then loaded on 10%-50% sucrose gradients poured using the Gradient Master (Serlabo Technologies) and centrifugated at 210.000 g for 2h35 at 4°C. 700 μl fractions were collected using the TELEDYNE ISCO collector while concomitantly acquiring corresponding 254nm absorbance.

### RNA extraction

#### RNA extraction from cells

Cells were harvested in 1ml of TRIzol reagent (Invitrogen) and total RNA was extracted according to manufacturer’s instructions. RNA concentration was assessed with a Nanodrop 2000/2000c spectrophotometer (ThermoScientific). 1 µg of RNA were used for reverse transcription assays using SuperScript II reverse Transcriptase Mix (Invitrogen) according to manufacturer’s instructions.

#### RNA extraction from sucrose gradients

50 pg of LUCIFERASE RNA (Promega) was added to 250 µl of fractions collected from sucrose gradients. 750 µl of TRIzol LS (Invitrogen) was then added and RNA was extracted according to the manufacturer’s instructions. The cDNAs were synthetized using SuperScript™ II Reverse Transcriptase (Invitrogen).

### Real Time qPCR assays

Quantitative PCR experiments were performed using SYBR Green Technology (Roche, Applied Biosystem) following the manufacturer’s instructions. Relative cDNA expression was normalized either by using mouse housekeeping gene encoding mRNAs β-Actin, Psmd9, Tbp and 603B20Rik (total cell RNA) or by using Luciferase mRNAs (sucrose gradient RNA). Serial dilutions were systematically performed to calculate qPCR efficiency, verify the qPCR linearity and determine normalized relative cDNA concentrations. Primers used in this study are listed in Table 6C.

### Metabolic labeling of protein synthesis

Cells were transfected with siRNAs in 6-well plates 48h before metabolic labelling. For labelling, the cells were incubated for 5 minutes at 37°C with 55µCi/well of 35S-L-methionine and ^35^S-L-cysteine Promix (Perkin Elmer) in a minimal volume of culture medium. To validate the labelling efficiency, a control was performed by incubating the cells with cycloheximide (100mg/mL final) for 10 min prior to labelling. After incubation, the cells were washed with 1 ml of ice-cold PBS and lysed in 500µL of RIPA buffer mixed with 2X final LDS Novex™ 4X Bolt™ loading buffer (ThermoFisher) for protein gel electrophoresis. Before loading onto precast Bis-Tris Bolt™ 4 to 12% acrylamide gels (ThermoFisher), the samples were sonicated for 5 min and denatured at 70°C for 10 min. The Simply Blue Safestain (Thermo) kit was used to check if protein loading was similar across lanes (accordingly to manufacturer’s guidelines). After coomassie staining, the gel was incubated in 30% ethanol, 10% acetic acid and 5% glycerol for 1h. The gel was dried at 75°C for 1h30 and autoradioactivity levels were then measured using the Typhoon Phosphor imager.

### Histone Immuno-histochemistry assays

Cells were centrifugated during 10 min at 377g and fixed in an alcohol based fixative solution Thinprep® (Hologic) during 15 min. After centrifugated during 5 min at 377g, the cells were resuspended in 10ml of Epredia™ Gel (Richard-Allan Scientific™ HistoGel™). The gel was hardened during 15 min at 4°C and the corresponding blocks were dehydrated and embedded in paraffin. 3µm-sections were immunostained using an antibody anti-histone H3 containing the trimethylated lysine 27 (H3K27me3) (Diagenode). Heat induced antigen retrieval was done using CC1 basic buffer (Ventana). Staining was performed using DAB Ultraview dection system (Ventana).

### RPL24 and RLP24 homologs protein alignments

The following protein sequences were considered for protein alignments. For RLP24 homologs: *S. cerevisiae* (Q07915), *C. elegans* RLP24 (Q17606), *D. rerio* RLP24 (Q7ZTZ2), *M. musculus* RSL24D1 (Q99L28), *R. norvegicus* RSL24D1 (Q6P6G7), *B. Taurus* RSL24D1 (Q3SZ12), *H. sapiens* RSL24D1 (Q9UHA3). For RL24 homologs: *S. cerevisiae* RL24A (P04449), *S. cerevisae* RL24B (P24000), *M. musculus* RL24 (Q8BP67). Multiple protein alignments were performed with the Clustal Omega software (https://www.ebi.ac.uk/Tools/msa/clustalo/) (82) and visualized with the Jalview software (http://www.jalview.org/) (83).

### RSL24D1 protein ternary structure predictions

The yeast RLP24 structures have been obtained from three yeast cryo-EM pre-60S structures available in the Protein Data Bank (https://www.rcsb.org): PDB-6N8J (residues 1-149, 2019, 3,50Å resolution) (45), PDB-6C0F (residues 2-130, 2018, 3,70Å resolution) (43) and PDB-3JCT (residues 1-150, 2016, 3,08Å resolution) (44). The mouse RSL24D1 protein structures have been modeled based on these 3 RLP24 cryo-EM protein structures using the SWISS-MODEL structures assessment tool (https://swissmodel.expasy.org/assess) (84-87) and respectively shared 61,48% (amino acids 1-135), 55,63% (amino acids 2-130) and 61,48% (amino acids 1-135) of homology with the yeast RLP24 protein sequence. The predicted mouse structures were compared with structural models of human RSL24D1 proteins derived from the PDB-6LSS and PDB-6LU8 pre-60S particle structures using the Pymol software.

### RNA-Seq sequencing and analysis

RNA libraries were prepared with the TruSeq Stranded Total-RNA kit and sequenced using an Illumina NovaSeq 6000 sequencing machine. Raw sequencing data quality controls were performed with FastQC (v 0.11.5). These data were aligned on the mouse genome (GRCm38) with STAR (v2.7.0f), with the annotation of known genes from gencode vM20, for careful quality control. RNA quality control metrics (library content, GC content) were computed using RSeQC (v 3.0.0) (88).

Gene expression was quantified using Salmon (0.14.1) on the raw sequencing reads, using the annotation of protein coding genes from gencode vM20 as index. Unless otherwise specified, the analyses were performed using R (v3.6.1). Starting from salmon transcript quantification, we used the R packages Tximport (v1.12.3) (89) DESeq2 (v1.24) (90) to perform the differential expression analyses (Wald test, and p-values correction with the Benjamini-Hochberg method).

### GO enrichment analysis

The analyses of overrepresented gene ontology categories were conducted using a reference set of 15093 genes expressed in CGR8 cells and using BiNGO (v3.0.3) (91) as well as the open source bioinformatics software platform Cytoscape (v3.7.2) (92) for visualization of the results. Only annotations with a corrected p-value < 0.01 were considered for further analysis (p-values corrections with the Benjamini-Hochberg method).

### Target genes analysis with StemChecker

Differentially expressed genes in Rsl24d1-depleted cells were analysed using the web-server StemChecker (http://stemchecker.sysbiolab.eu/) (51), without masking the cell proliferation and cell cycle genes.

### Analysis of binding sites defined by ChIP-seq

To identify binding sites for TFs and chromatin-associated factors in the ±1 kb region of TSSs, we combined all datasets available in the ChIP-Atlas database (52) corresponding to mouse wild type ESCs and mouse differentiated cell types for each factor. To select binding sites preferentially bound by the selected factors in ESCs, we selected all sites with a ratio of ESC average score / Differentiated averaged score >2.

## Acknowledgments

We thank Drs. P. Savatier and P.Y. Bourrillot (Stem cell and Brain Research Institute, Lyon, France) for providing protein extracts of human ESCs, and Pr. A. Nagy (University of Toronto, Canada) for providing the R1 and G4 cell lines. We thank R. Pommier and A. Viari from the Gilles Thomas Bioinformatics Platform (CRCL, Lyon, France) for the analysis of RNAseq data. This study was supported in part by a Marie Curie Career Integration Grant 631794 from the European Union and by a national ATIP/AVENIR grant from the Institut National de la Santé et de la Recherche Médicale (INSERM) to M.G., and by a Ph.D. fellowship by La Ligue Nationale Contre le Cancer (M.B.), by grants from Labellisation de la ligue contre le cancer, Agence Nationale pour la Recherche (ANR) and Institut National du Cancer (INCa) (A.H. and F.L.).

## Author contributions

S.D., M.B. and M.G. designed the research. S.D., M.B., F.B., B.B., J.B., C.I., A.H., D.M. performed the research. S.D., M.B., C.V., F.C., J.J.D., F.L, E.R., F.D. and M.G. analyzed the data. S.D., M.B., F.B. and M.G. wrote the manuscript. All authors discussed the results and commented on the manuscript.

