## Supplementary figures and images for "RSL24D1 sustains steady-state ribosome biogenesis and pluripotency translational programs in embryonic stem cells"

### Supplemental Figure 1

Figure Sup. 1, Durand et al.

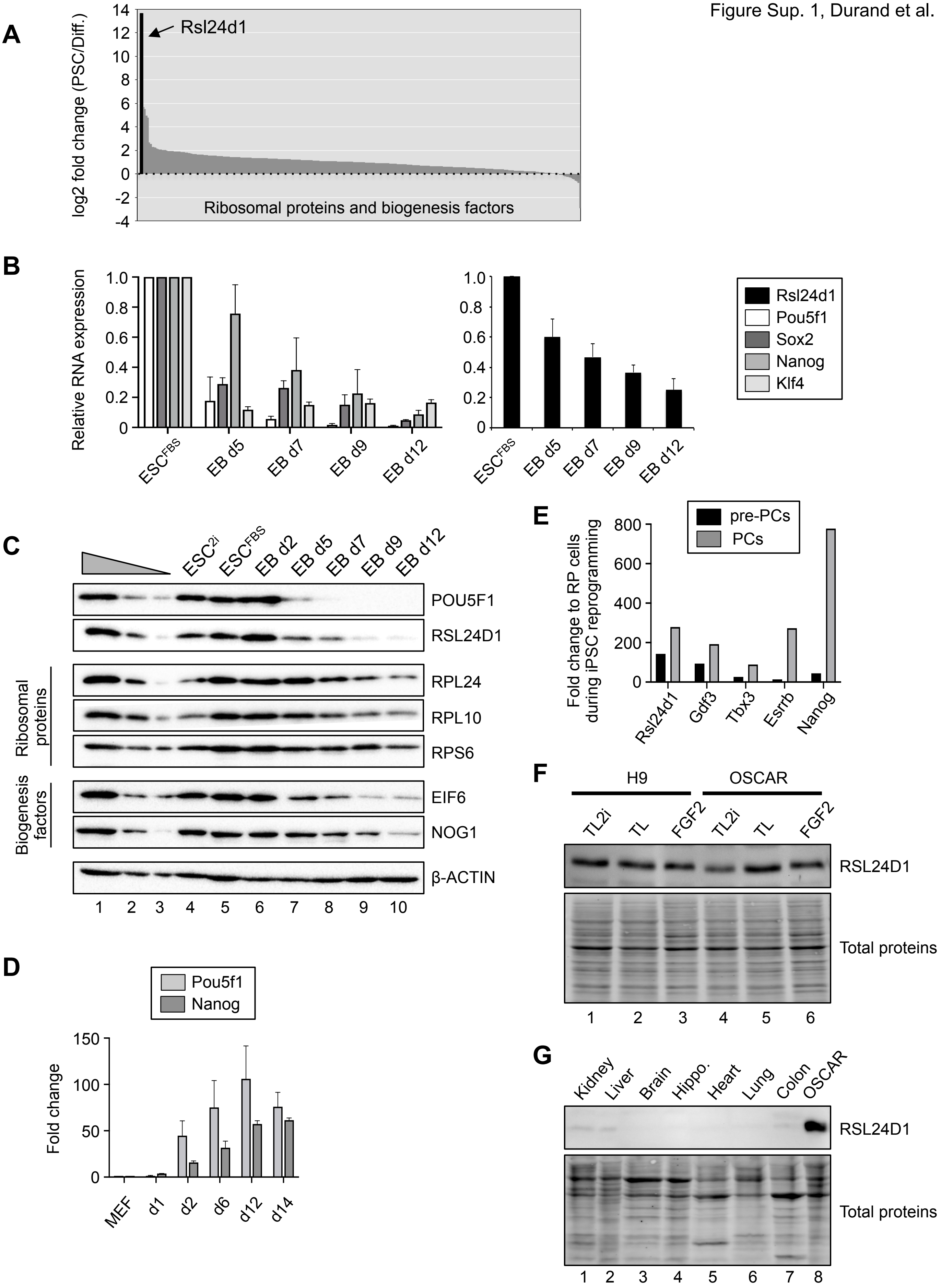

### Supplemental Figure 2

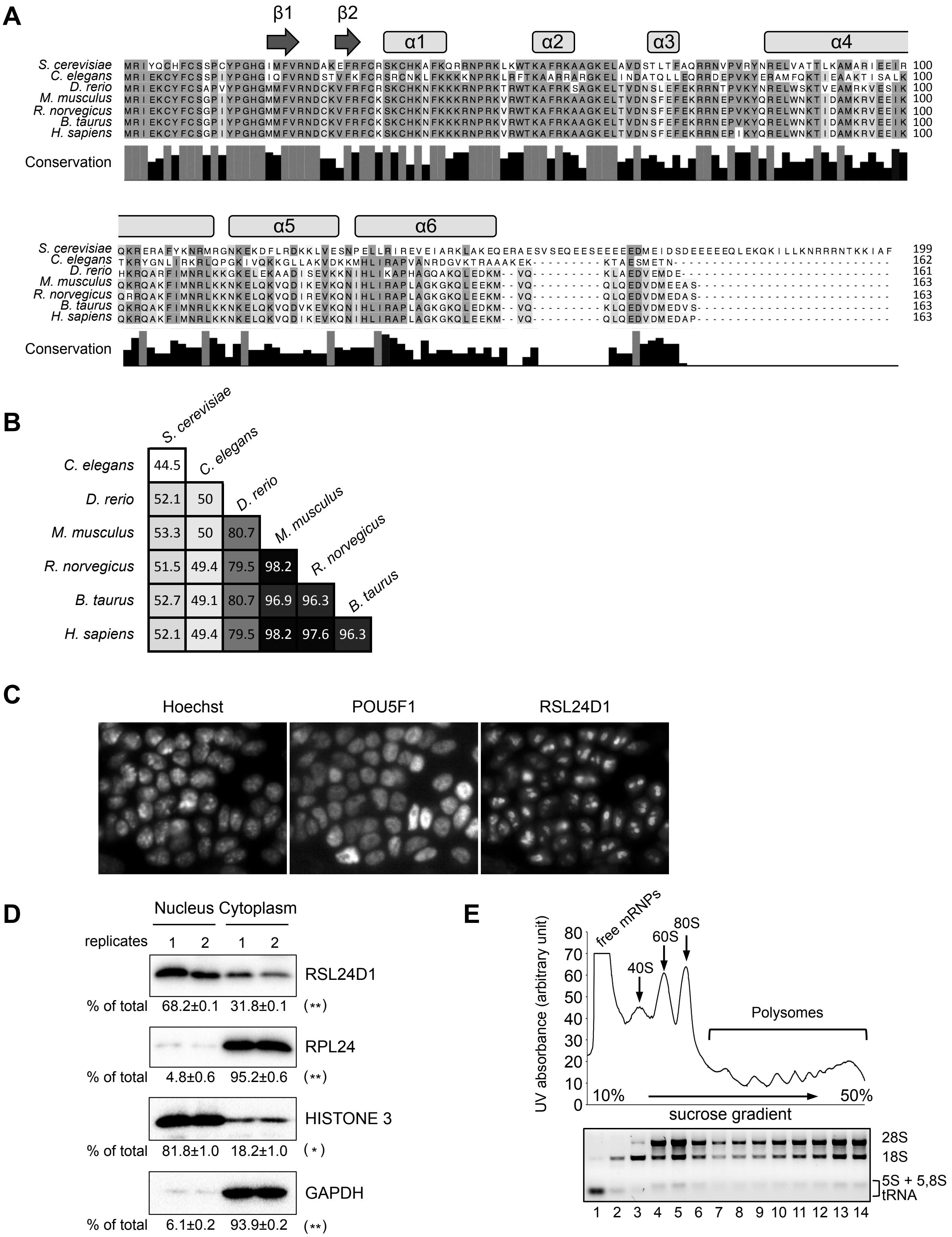

### Supplemental Figure 3

Figure Sup. 3, Durand et al.

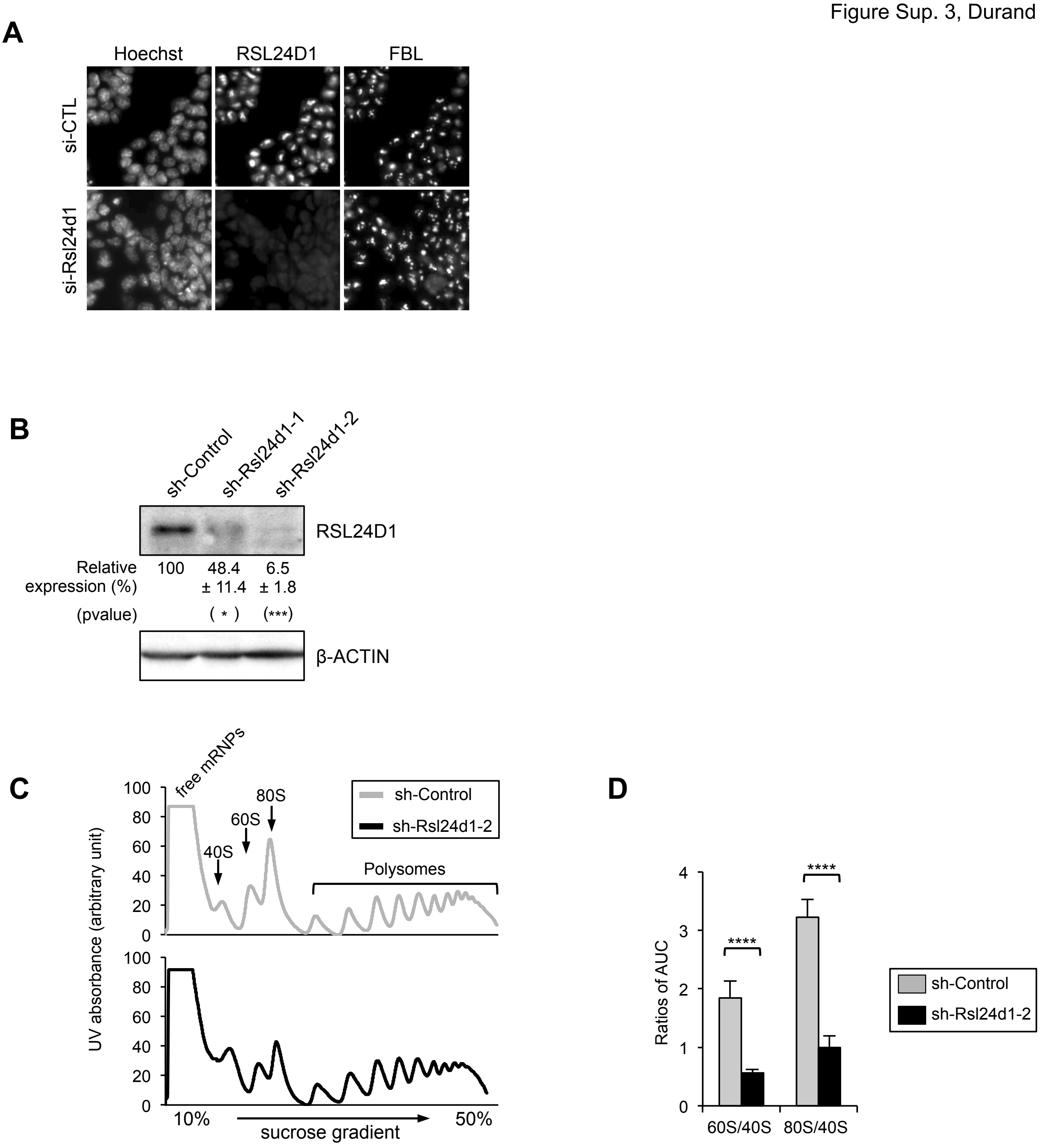

### Supplemental Figure 4

Figure Sup. 4, Durand et al.

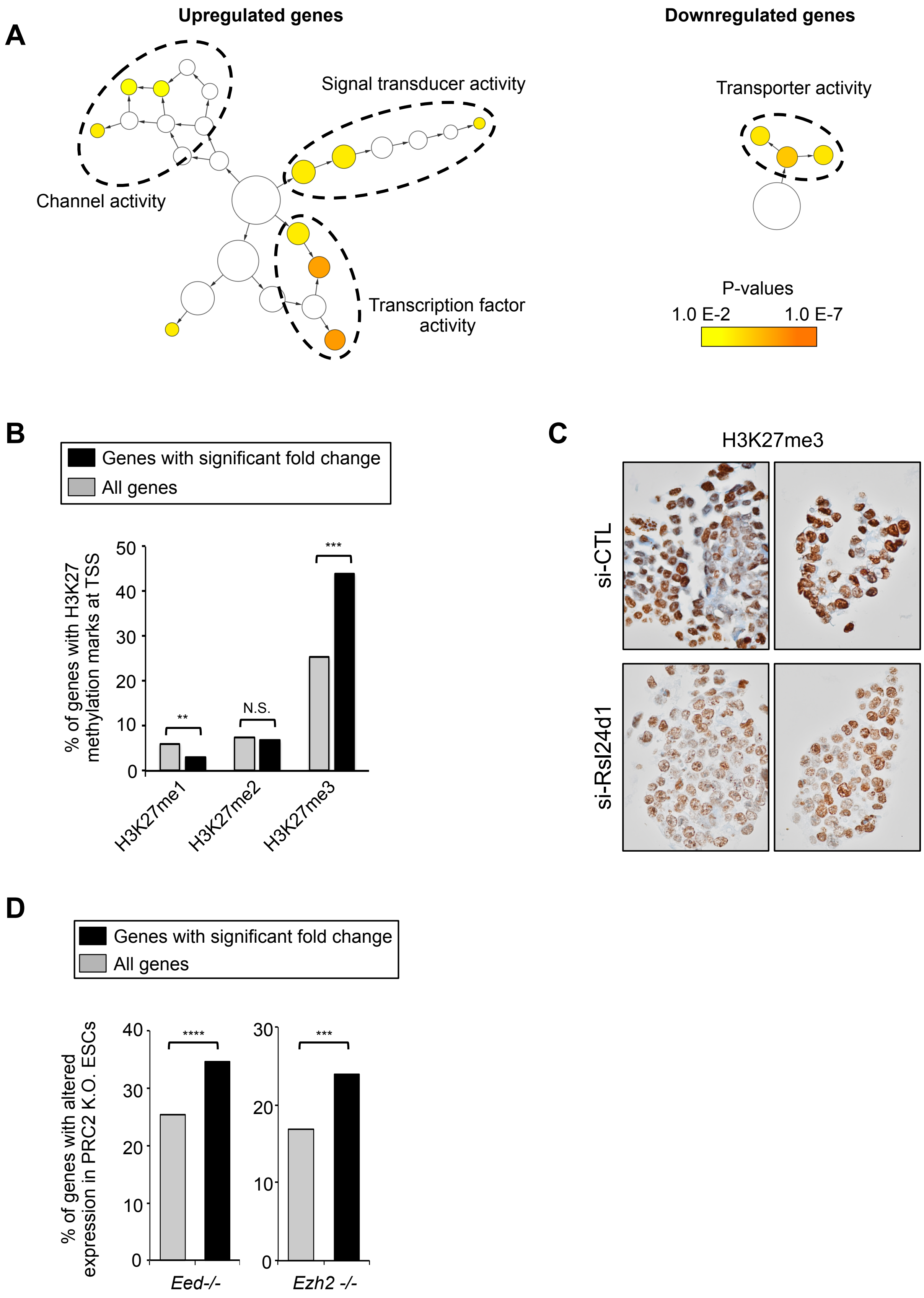

### Supplemental Figure 5

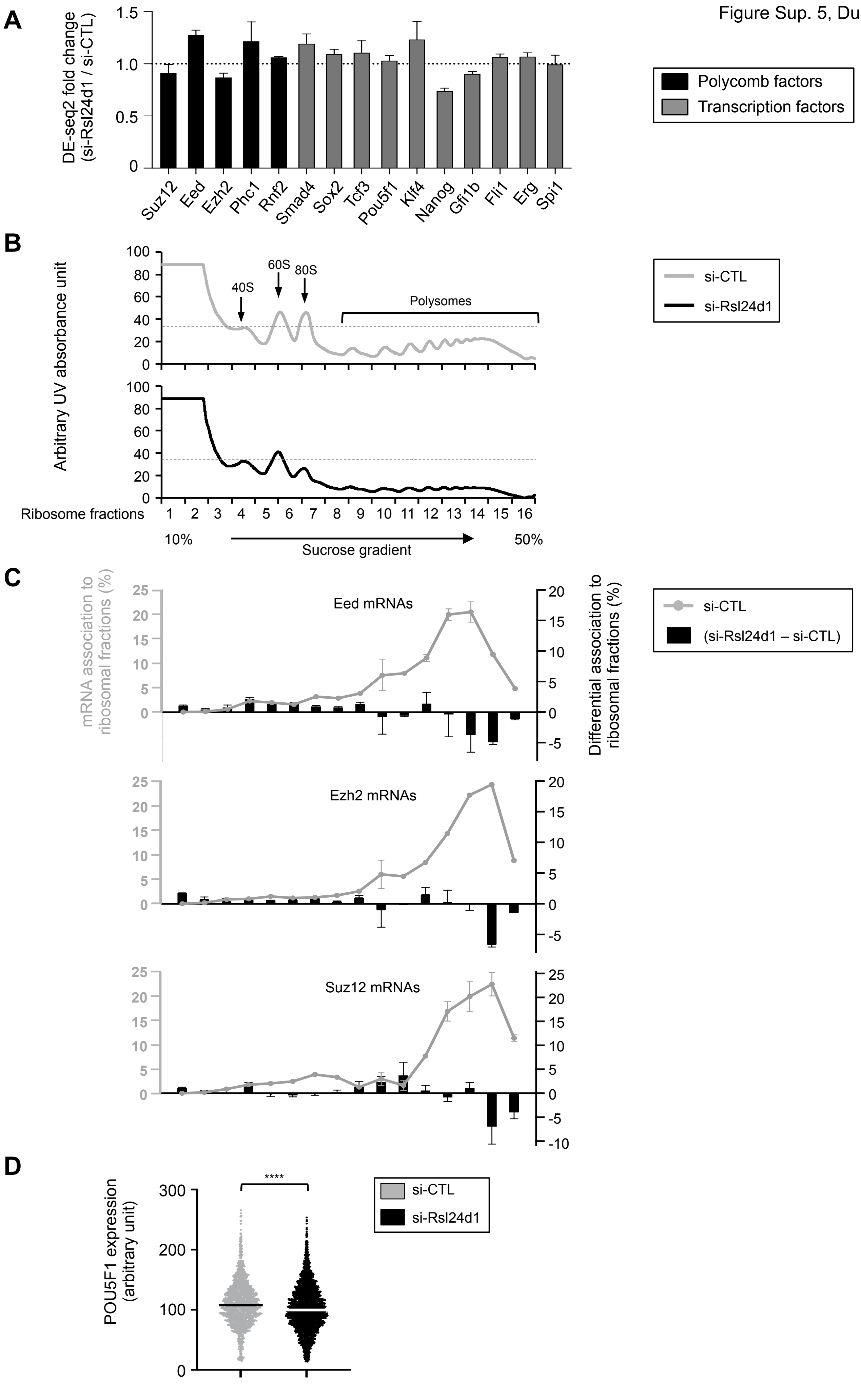

### Supplemental Figure 6

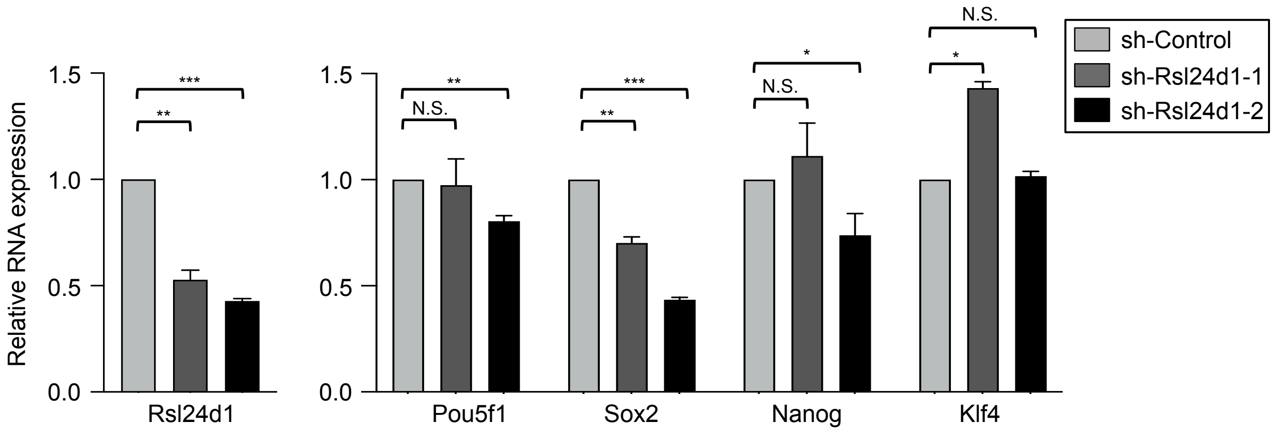

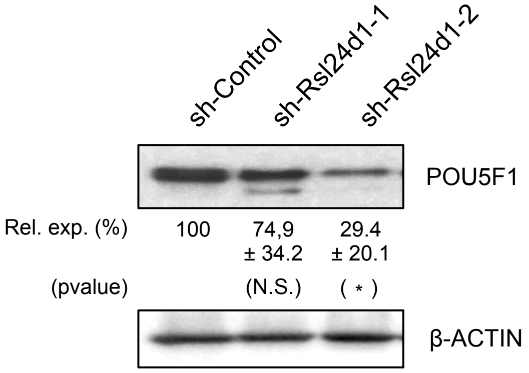

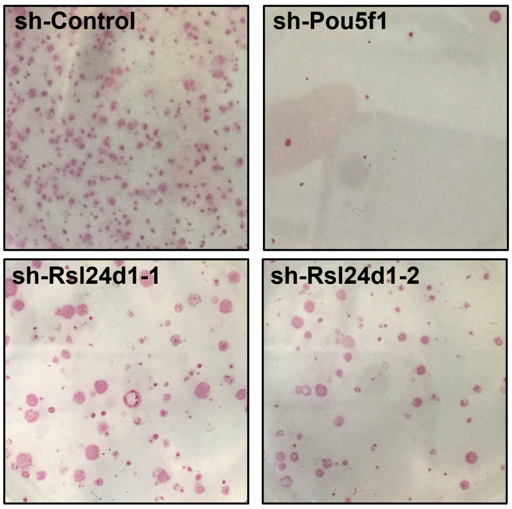
