## Supplemental Figure Legends for "RSL24D1 sustains steady-state ribosome biogenesis and pluripotency translational programs in embryonic stem cells"

**Supplementary Figure Legends**

**Supplemental Figure 1. RSL24D1 expression pattern in murine and human pluripotent cells.**

(A) RNA-seq profiling of expression changes for 303 mRNAs encoding ribosomal proteins and ribosome biogenesis factors between mouse pluripotent and differentiated cells detailed in Table S1. Expression changes are shown as log2 ratios of the average expression in PSCs (n= 7 cell lines) over differentiated cells (n= 6 cell lines), and ranked from the most enriched mRNAs in PSCs (left) to the most enriched in differentiated samples (right).

(B) Quantitative PCR analysis of the expression of Pou5f1, Sox2, Nanog and Klf4 (left panel) and Rsl24d1 mRNAs (right panel) during a kinetic of CGR8 differentiation into embryoid bodies (EBs) at days 0 (ESC^FBS^), 5, 7, 9 and 12 (n=3). Transcript levels are normalized to the β-Actin and Psmd9 mRNAs.

(C) Representative immunoblots of RSL24D1, POU5F1, RPL24, RPL10 RPS6, EIF6 and NOG1 in total extracts from ground state (ESC^2i^) and naïve pluripotent CGR8 (ESC^FBS^) as well as EBs at 2, 5, 7, 9 and 12 days of differentiation. Lanes 1 to 3 correspond to serial dilutions of ESC^FBS^ extracts (1:1, 1:3 and 1:9, respectively). β-ACTIN is shown as a loading control.
(D) Quantitative PCR analysis of the kinetics of expression of endogenous Pou5f1 and Nanog mRNAs during the course of reprogramming of MEFs derived from doxycycline-inducible Col1a1-tetO-OKMS mice into iPSCs. Transcript levels are indicated relative to MEFs (day 0), at day 1, 2, 6, 12 and 14 post doxycycline induction, and are normalized to the Tbp and Psmd9 mRNAs.

(E) Single-cell RNA-seq gene expression changes in cells undergoing somatic reprogramming (pre-PCs) and chimera-competent pluripotent cells (PCs) compared to cells with reprogramming potential (RP), as described by Guo and colleagues (40).

(F) Representative immunoblot of RSL24D1 in total extracts from H9 and OSCAR human ESC lines maintained in naïve-like conditions (TL2i), in primed conditions with LIF + 4’OHT supplemented media (TL) supporting STAT3 overexpression or in presence of FGF2, as described in (41). TCE labeling of tryptophan-containing proteins (referred as Total proteins) is used as loading control.

(G) As described in (F) in total extracts from a panel of human adult tissues and from pluripotent human OSCAR ESCs cultured in the presence of ectopic FGF2.

**Supplemental Figure 2. RSL24D1 sequence and structure are conserved across evolution.**

(A) Multiple protein alignments of yeast Rlp24 with homologs from higher eukaryote species. Amino acid numbers are indicated on the right for each homolog protein, and the conservation score to the yeast Rlp24 sequence for each position is calculated by the Jalview software and indicated below. The secondary structure predictions for the first 149 amino acids of the yeast Rlp24 protein are indicated above. β-shits are represented by arrows and α-helixes by boxes.
(B) Amino acid sequence identity calculated by paired-alignment between yeast Rlp24 and homologs from higher eukaryote species. RSL24D1 is highly conserved in higher eukaryotes from zebrafish to human.

(C) Representative images of naïve CGR8 cells in ESC^FBS^ conditions stained with Hoechst and with anti-OCT4 and anti-RSL24D1 antibodies (20X objective).

(D) Nucleo-cytoplasmic fractionation in two independent biological replicates of CGR8 cells cultured in naive conditions (ESC^FBS^). Representative western blots of RSL24D1 and RPL24. Immunoblots of HISTONE H3 and GAPDH are shown as specific nuclear and cytoplasmic proteins, respectively. Quantifications of RSL24D1 and RPL24 signals in both nuclear and cytoplasmic fractions are estimated as percentage of total signals, defined by the sum of nuclear and cytoplasmic signal. ** pval<0.01, * pval<0.05, paired two-tailed Student’s t test.
(E) related to Figure 3D. Polysome profiling by centrifugation on sucrose gradient of CGR8 ESC^FBS^ cytoplasmic extracts. Free mRNPs, 40S, 60S, 80S monosomes and polysomes are detected by UV-absorbance and indicated on the absorbance curve. Total RNAs were extracted from collected fractions, analyzed on non-denaturing agarose gels and revealed with ethidium bromide. The positions of the 28S, 18S, 5.8S, 5S rRNAs and tRNAs are indicated.

**Supplemental Figure 3. RSL24D1 depletion alters ribosome biogenesis and protein translation.**
(A) Representative images of si-CTL and si-Rsl24d1 treated naïve CGR8 cells stained with Hoechst and with anti-FBL and anti-RSL24D1 antibodies (20x objective).

(B) Representative immunoblot of RSL24D1 (n=3) in total protein lysates from naïve CGR8 cells stably expressing control shRNAs or 2 independent Rsl24d1-targeting shRNAs. β-ACTIN is used as a loading control. RSL24D1 signals are normalized to β-ACTIN signals and expressed relative to the control shRNA expressing cells. *** pval<0.005, * pval<0.05, unpaired two-tailed Student’s t test.

(C) Representative polysome profiling performed by centrifugation on sucrose gradients of cytoplasmic extracts from naive CGR8 cells infected with lentiviruses expressing control non-targeting shRNAs (grey) or shRNAs targeting Rsl24d1 (sh-Rsl24d1-2, black). 40S, 60S, 80S monosomes and polysomes are detected by UV-absorbance and indicated on the absorbance curve.
(D) Histogram indicating ratios of 60S/40S and 80S/40S absorbance peaks calculated by determining the area under the curve (AUC) for the 40S, 60S and 80S absorbance signals (n=5). **** pval<0.0001, unpaired two-tailed Student’s t test.

**Supplemental Figure 4. RSL24D1 depletion impairs specific molecular pathways.**
(A) Gene ontology enrichment analyses of molecular function annotation terms for genes displaying significant expression changes in RNA-seq analysis of CGR8 cells treated with Rsl24d1 siRNAs. The left and right panels represent hierarchical trees of the most enriched terms in genes upregulated and downregulated in Rsl24d1-depleted cells, respectively. The size of the nodes represents the numbers of genes associated to each GO term, and the corresponding p-values are indicated by colour codes, according to the scale provided. The most represented GO term categories are indicated. The corresponding data are available in Table S3.
(B) Analysis of the proportion of transcription start sites (TSS) associated with H3K27 mono, bi or tri-methylation marks (9) for genes differentially expressed in Rsl24d1-depleted CGR8 cells (n=529, black) and in the entire set of genes associated with significant expression predictions (DEseq2 corrected p values < 0.01) from the RNA-seq analysis, regardless of the change in expression (n=3111, grey). *** (pval<0.0005), ** (pval<0.005), N.S.: not significant (pval>0.05), Fisher exact test.

(C) Representative images of immunohistochemistry assays stained with anti trimethylated form of H3K27 (H3K27me3) antibodies, in CGR8 colonies cultured in ESC^FBS^ conditions and treated with si-CTL or si-Rsl24d1 siRNAs.

(D) Analysis of the proportion of genes showing altered expression in *Eed*-/- (7) or *Ezh2*-/- (8) mouse ESCs, respectively, that overlaps with the 529 genes associated with a differential expression in si-Rsl24d1-treated naive CGR8 cells (black) and in the entire set of 3111 genes associated with a significant expression predictions (DEseq2 corrected pval<0,01), as described in (B) (grey). **** pval<0.0001, *** pval<0.001, Fisher exact test.

**Supplemental Figure 5. Rsl24d1 downregulation alters the translation of PRC2-component mRNAs and POU5F1 levels in CGR8 cells.**

(A) Bar graph representing the ratio of expression estimated by DE-seq2 in si-Rsl24d1 treated CGR8 cells over si-Ctl treated cells, for PRC factors (black) and PTFs (grey) analyzed in Figure 4B.
(B) Polysome profiling obtained after centrifugation on sucrose gradients of cytoplasmic extracts from naïve pluripotent CGR8 cells treated with non-targeting (si-CTL, grey) or Rsl24d1-targeting siRNAs (si-Rsl24d1, black). 40S, 60S, 80S monosomes and polysomes are detected by UV-absorbance and indicated on the absorbance curve.

(C) Graphs representing Eed, Ezh2 and Suz12 mRNA levels measured by RT-qPCR in each fraction collected from the polysome profiling (grey curves, left scales) after normalization with a Luciferase spike-in mRNA and relative to the total amount of mRNAs detected in the polysome profiling (sum of all fractions) (n=3). The bar graph represents the differential detection of each mRNA in each fraction between CGR8 cells treated with si-Rsl24d1 and control siRNAs (black bars, right scales).

(D) Quantifications of immunostaining signals from siRNA treated naive CGR8 cells stained with anti-POU5F1 antibodies and imaged by the Operetta CLS high-content analysis microscope (si-CTL, n=1620; si-Rsl24d1, n=2740). Mean values are indicated by a black (si-CTL) and a white (si-Rsl24d1) line in the scatter plots. **** pval<0.0001, unpaired two-tailed Student’s t test.

**Supplemental Figure 6. RSL24D1 expression is required to sustain ESC self-renewal.**
(A) RT-qPCR analyses of Rsl24d1, Pou5f1, Sox2, Nanog and Klf4 mRNAs in CGR8 cells expressing non-targeting shRNAs or 2 independent shRNAs targeting Rsl24d1. Transcript levels are normalized to the β-Actin and Tbp mRNAs and expression changes are indicated as fold change relative to sh-control treated cells (n=3). *** pval<0.0001, ** pval<0.005, * pval<0.05, N.S.: not significant (pval>0.05), paired two-tailed Student’s t test.

(B) Representative immunoblot of POU5F1 in naive CGR8 cells treated with non-targeting or Rsl24d1-targeting shRNAs. β-ACTIN signals are used for normalization of protein loading and immunodetection. Quantifications of the POU5F1 signals normalized to β-ACTIN signals and relative to the (Rel. exp. (%)) are indicated below. The indicated P values are relative to the non-targeting shRNA expressing conditions (unpaired two-tailed Student’s t test): * (pval<0.05), N.S.: not significant (pval>0.05).

(C) Data related to Figure 6C. Representative images of alkaline phosphatase stainings detected in colonies formed by naive CGR8 expressing non-targeting shRNAs, Pou5f1-targeting shRNAs, or two independent shRNAs targeting Rsl24d1.
